# Phosphoglycerate kinase 1 cooperates with AP-2 complex to promote endocytosis

**DOI:** 10.1101/2025.10.29.685329

**Authors:** Shao-Ling Chu, Yu-Tzu Chang, Che-Yu Hung, Chia-Jung Yu, Jia-Wei Hsu

## Abstract

Adaptor proteins play a critical role in intracellular protein transport by binding to cargoes and facilitating their sorting into vesicles. The adaptor protein 2 (AP-2) complex specifically recognizes sorting motifs on cargo proteins such as epidermal growth factor receptor (EGFR) and transferrin receptor (TfR) for proper cargo sorting. However, the coordination of these adaptor proteins in cargo recognition remains unclear. Here, we uncover the non-canonical role of glycolytic enzyme phosphoglycerate kinase 1 (PGK1) in protein endocytosis, acting independently of its catalytic activity. PGK1 serves as a co-adaptor to directly engage cargoes and reinforce AP- 2 complex stability, orchestrating efficient cargo sorting to promote endocytosis. Furthermore, PGK1 also integrates into the AP-2-clathrin complex, enhancing cargo recognition and internalization. Collectively, our findings establish a previously unidentified coat protein complex containing PGK1, highlighting its dual role as a cargo adaptor and stabilization of the AP-2 complex.

## INTRODUCTION

Coat proteins play a central role in intracellular protein transport by binding to cargoes and facilitating their sorting into specific vesicles ^1,2^. Cargo proteins interact directly with different coat proteins, establishing their dynamic movement between organelle compartments within the secretory and endocytic pathways ^3^. The best characterized coat protein for controlling endocytosis is the clathrin-adaptor protein 2 (AP-2) coat complex ^4^. The clathrin triskelion couples with the AP-2 complex to form the clathrin-AP-2 coat complex, which functions in protein endocytosis ^5–7^. The AP-2 adaptor protein complex consists of two large subunits (α and β), a medium subunit (µ), and a small subunit (σ). Two of the best characterized sorting motifs on cargo proteins, the acidic dileucine motif and tyrosine-based motif on epidermal growth factor receptor (EGFR) and the tyrosine-based motif on transferrin receptor (TfR), can be specifically recognized by the AP-2 complex ^8^. The acidic dileucine motif binds to AP-2σ, whereas the tyrosine-based motif interacts with AP-2µ ^9,10^. Upon being recruited to the plasma membrane, the AP-2 complex recognizes sorting signals on cargo proteins and subsequently recruits clathrin to facilitate vesicle formation. During this process, multiple accessory proteins interact with the AP-2 complex directly or indirectly to form the clathrin-AP-2 coat complex ^8^. These proteins, with the AP-2 complex as the central component, form an interconnected system to regulate the dynamics of protein endocytosis. However, how these adaptor proteins coordinate to specifically recognize cargo proteins remains unclear.

Phosphoglycerate kinase 1 (PGK1) is a metabolic enzyme that catalyses adenosine triphosphate (ATP) production during glycolysis. In addition to its canonical role in glycolysis, PGK1 functions as a protein kinase in tumorigenesis and autophagy ^11,12^. By further investigating endocytic protein transport, we previously identified that PGK1 directly interacts with the dileucine sorting motif on EGFR to promote its lysosomal protein transport ^13^. We also found that the ability of PGK1 to promote EGFR lysosomal transport is independent of its kinase activity, indicating that its moonlighting function does not require its metabolic activity ^13^. Here, we demonstrate that PGK1 unexpectedly has a moonlighting function in regulating protein endocytosis. We show that the PGK1 and AP-2 complex specifically recognize sorting motifs on cargo proteins to increase binding avidity via multivalent interactions and further recruit clathrin to account for endocytosis. These mechanisms provide functional parallels of PGK1 to the lysosomal protein transport pathway. Overall, our study revealed a clathrin protein complex containing PGK1 as the cargo adaptor in cooperation with AP-2 complex for proper cargo sorting during protein endocytosis.

## RESULTS

### PGK1 promotes AP-2-dependent endocytosis

We previously reported that PGK1 acts as a cargo adaptor to promote the lysosomal transport of EGFR upon EGF stimulation ^13^. To understand the additional role of PGK1 in endocytic transport, we first performed coimmunoprecipitation-coupled mass spectrometry to identify PGK1-interacting proteins on the endosomal membrane. We identified one potential target, adaptor protein 2 complex alpha (AP-2α), from the top hit of our proteomic analysis (Table EV1). The AP-2 complex regulates major clathrin-dependent endocytosis events, including the endocytosis of many cellular receptors, such as transferrin receptor (TfR) and epidermal growth factor receptor (EGFR). Interference with the AP-2 component significantly impairs the endocytosis of TfR and EGFR ^14^. Notably however, a previous report demonstrated that some endocytosis events still occur in siAP-2-treated cells ^15^, leading us to consider additional or alternative factor(s) that can cooperate with the AP-2 complex to control endocytosis. We hypothesized that PGK1 may cooperate with the AP-2 complex to participate in AP-2-dependent endocytosis. We first examined TfR endocytosis in cells by using flow cytometry to detect internalized fluorescein-labelled Tf signals combined with the use of Trypan blue to quench plasma membrane-bound forms ^16,17^. We then found that single knockdown of AP-2α or PGK1 significantly impaired TfR endocytosis, and co-knockdown of AP-2α and PGK1 showed synergistic effect on the inhibition of endocytosis (Fig. 1A). We next addressed the kinetic analysis of endocytosis by tracking the cargo internalization of Tf-labelled TfRs into early endosomes combined with acid wash to remove pre-bound ligands on the cell surface, and also revealed a significant defect in TfR endocytosis in AP-2α and PGK1 co-knockdown cells (Fig. 1B, 1C, and EV1A, EV1B). The exacerbation of EGFR endocytosis defects was also detected in AP-2α and PGK1 co-knockdown cells compared with single knockdown cells (Fig. 1D, 1E, and EV1C). In addition, we also confirmed the involvement of the mu subunit of the AP-2 complex (AP-2μ) to cause the severe defect in AP-2-dependent endocytosis by tracking the internalization of TfR and EGFR in cells in which both AP-2μ and PGK1 were largely depleted (Fig. 1B-1E and EV1B-1C). As a control, we also showed that the internalization of the cholera toxin B subunit (CTxB) was not affected in AP-2α and PGK1 co-knockdown cells (Fig. EV1D). To avoid the effects of low temperature during ligand prebinding, we simply treated the ligand at 37°C to track receptor endocytosis ^17^. Consistently, we also observed that the uptake of TfR was significantly abolished in AP-2α and PGK1 co-knockdown cells compared with that in siPGK1 or siAP-2α-treated cells (Fig. EV1E). These results demonstrated that PGK1 and AP-2α synergistically cause endocytosis defects. We further determined that the glycolytic activity of PGK1 was not required for AP-2- dependent endocytosis because both the wild-type PGK1 and the catalytic mutant PGK1-T378P successfully restored TfR endocytosis in siPGK1-treated cells (Fig. 1F, 1G, and EV1F), and for the restoration of EGFR endocytosis in cells subjected to PGK1 knockdown was also observed (Fig. 1H, 1I, and EV1G). Together, these data suggest that PGK1 can be the essential protein factor to regulate protein endocytosis independent of its catalytic activity, especially when the function of the AP-2 complex is impaired.

**Figure 1.**
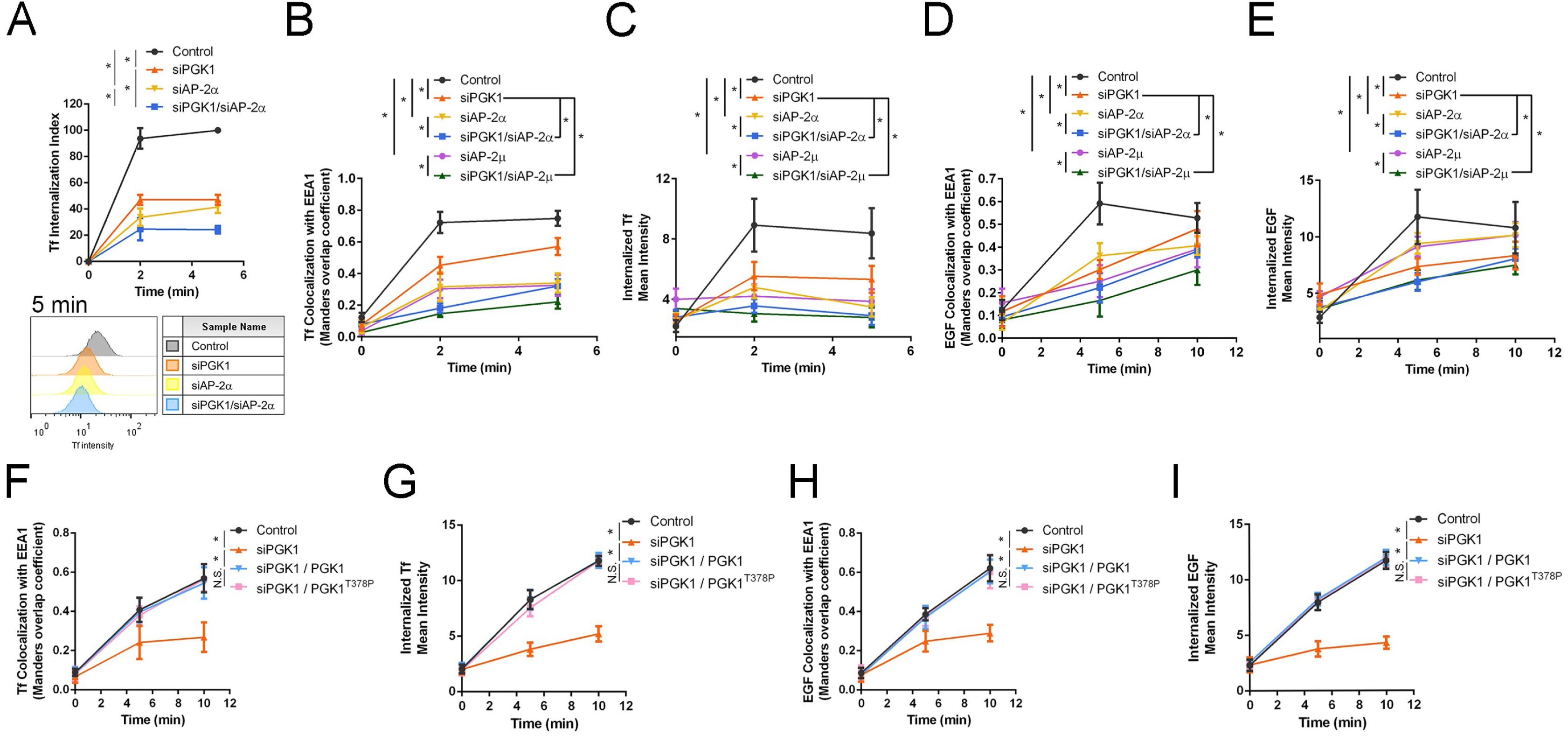
PGK1 regulates AP-2-mediated endocytosis independently of its kinase activity. Quantitative analyses of cells treated with the indicated siRNAs are presented as means ± standard deviations (SDs). \**P*<0.01; NS *P*>0.05 **(A)** Internalization of Tf was quantified by flow cytometry using trypan blue quenching at 0, 2, and 5 minutes (n>10,000 cells, four independent experiments). One-way ANOVA with Tukey’s test at the 5-minute time point: control vs. siPGK1 (*P*=0.0003); control vs. siAP-2α (*P*=0.0004); siAP-2α vs. siPGK1/siAP-2α (*P*=0.0253); siPGK1 vs. siPGK1/siAP-2α (*P*=0.0081). **(B)** Colocalization of Tf with EEA1, expressed as the Manders’ overlap coefficient, was quantified at 0, 2, and 5 minutes (n=20 cells, two independent experiments). One-way ANOVA with Tukey’s test at 2-minute time point: control vs. siPGK1 (*P*=2.4×10^-14^); control vs. siAP-2α (*P*=2.4×10^-14^); siPGK1 vs. siPGK1/siAP-2α (*P*=3.8×10^-10^); siAP-2α vs. siPGK1/siAP-2α (*P*=8.3×10^-10^); control vs. siAP-2μ (*P*=2.4×10^-14^); siPGK1 vs. siPGK1/siAP-2μ (*P*=2.4×10^-14^); siAP-2μ vs. siPGK1/siAP-2μ (*P*=1.8×10^-12^). **(C)** Mean intensity of internalized Tf was assessed at 0, 2, and 5 minutes (n=20 cells, two independent experiments). One-way ANOVA with Tukey’s test at the 2-minute time point: control vs. siPGK1 (*P*=5.1×10^-14^); control vs. siAP-2α (*P*=2.5×10^-14^); siPGK1 vs. siPGK1/siAP-2α (*P*=3.1×10^-8^); siAP-2α vs. siPGK1/siAP-2α (*P*=0.0013); control vs. siAP-2μ (*P*=2.4×10^-14^); siPGK1 vs. siPGK1/siAP-2μ (*P*=5.2×10^-12^); siAP-2μ vs. siPGK1/siAP-2μ (*P=*0.0028). **(D)** Colocalization of EGF with EEA1, expressed as the Manders’ coefficient, was quantified at 0, 5, and 10 minutes (n=20 cells, two independent experiments). One-way ANOVA with Tukey’s test at the 5-minute time point: control vs. siPGK1 (*P*2.5×10^-14^); control vs. siAP-2α (*P*=6.3×10^-14^); siPGK1 vs. siPGK1/siAP-2α (*P*=0.0044); siAP-2α vs. siPGK1/siAP-2α (*P*=2.3×10^-8^); control vs. siAP-2μ (*P*=2.4×10^-14^); siPGK1 vs. siPGK1/siAP-2μ (*P*=6.1×10^-8^); siAP-2μ vs. siPGK1/siAP-2μ (*P*=0.0016). **(E)** Mean intensity of internalized EGF was assessed at 0, 5, and 10 minutes (n=25 cells, two independent experiments). One-way ANOVA with Tukey’s test at the 5-minute time point: control vs. siPGK1 (*P≈*0); control vs. siAP-2α (*P*=7.9×10^-9^); siPGK1 vs. siPGK1/siAP-2α (*P*=0.0024); siAP-2α vs. siPGK1/siAP-2α (*P≈*0); control vs. siAP-2μ (*P*=9.7×10^-11^); siPGK1 vs. siPGK1/siAP-2μ (*P*=0.0129); siAP-2μ vs. siPGK1/siAP-2μ (*P*=7.6×10^-13^). **(F)** Colocalization of Tf with EEA1, expressed as Manders’ coefficient, was performed to compare the effects of PGK1 and PGK1^T378P^ at 0, 5, and 10 minutes (n=15 cells, two independent experiments). One-way ANOVA with Tukey’s test at the 5-minute time point: control vs. siPGK1 (*P*=5.9×10^-8^); siPGK1 vs. PGK1 (*P*=3.4×10^-7^); PGK1 vs. PGK1^T378P^ (*P*=0.9055). **(G)** Mean intensity of internalized Tf was measured to compare the effects of PGK1 and PGK1^T378P^ at 0, 5, and 10 minutes (n=15 cells, two independent experiments). One-way ANOVA with Tukey’s test at the 5-minute time point: control vs. siPGK1 (*P*=7.3×10^-12^); siPGK1 vs. PGK1 (*P*=7.3×10^-12^); PGK1 vs. PGK1^T378P^ (*P*=0.0698). **(H)** Colocalization of EGF with EEA1, expressed as Manders’ coefficient, was performed to compare the effects of PGK1 and PGK1^T378P^ at 0, 5, and 10 minutes (n=15 cells, two independent experiments). One-way ANOVA with Tukey’s test at 5-minute time point: control vs. siPGK1 (*P*=1.6×10^-9^); siPGK1 vs. PGK1 (*P*=4.4×10^-8^); PGK1 vs. PGK1^T378P^ (*P*=0.9835). **(I)** Mean intensity of internalized EGF was performed to compare the effects of PGK1 and PGK1^T378P^ at 0, 5, and 10 minutes (n=15 cells, two independent experiments). One-way ANOVA with Tukey’s test at the 5-minute time point: control vs. siPGK1 (*P*=7.3×10^-12^); siPGK1 vs. PGK1 (*P*=7.3×10^-12^); PGK1 vs. PGK1^T378P^ (*P*=0.9728).

### PGK1 interacts with AP-2α and stabilizes the AP-2 complex

We next investigated the association of PGK1 with the AP-2 complex and revealed that PGK1 interacts with the AP-2α and AP-2σ subunits (Fig. 2A). We further confirmed that PGK1 associated with the endogenous α and μ subunits of the AP-2 complex (Fig. 2B). A proximity ligation assay (PLA) also demonstrated the interaction of PGK1 and AP-2α in intact cells (Fig. 2C and 2D). Interestingly, the PGK1 and AP-2α interaction was largely detected during TfR and EGFR endocytosis and peaked at the initial stage after the induction of endocytosis (Fig. 2C-2D and EV2A-2B). These data suggest that PGK1 associates with the AP-2 complex upon the initiation of endocytosis. We next investigated the possibility of a direct interaction between PGK1 and the AP-2 complex and found that recombinant PGK1 and AP-2α directly interact with one another (Fig. 2E). We next postulated that AP-2α can serve as a bridge to link PGK1 with other subunits of the AP-2 complex. Although PGK1 weakly interacts with the μ and σ subunits of the AP-2 complex, in the presence of AP-2α, the binding of AP-2μ and AP-2σ to PGK1 was noticeably stronger (Fig. 2F and 2G), suggesting that the AP-2α subunit serves as a linker in the binding of PGK1 and the AP-2 complex. This interaction was further confirmed in cells by showing the reduction in PGK1 interactions with the AP-2β subunit in siAP-2α-treated cells (Fig. 2H). However, the association of the AP-2 complex was not dramatically affected in siPGK1- treated cells (Fig. EV2C). Collectively, these data suggest that AP-2α plays a central role in the binding of PGK1 and the AP-2 subunits during endocytosis (Fig. EV2D).

**Figure 2.**
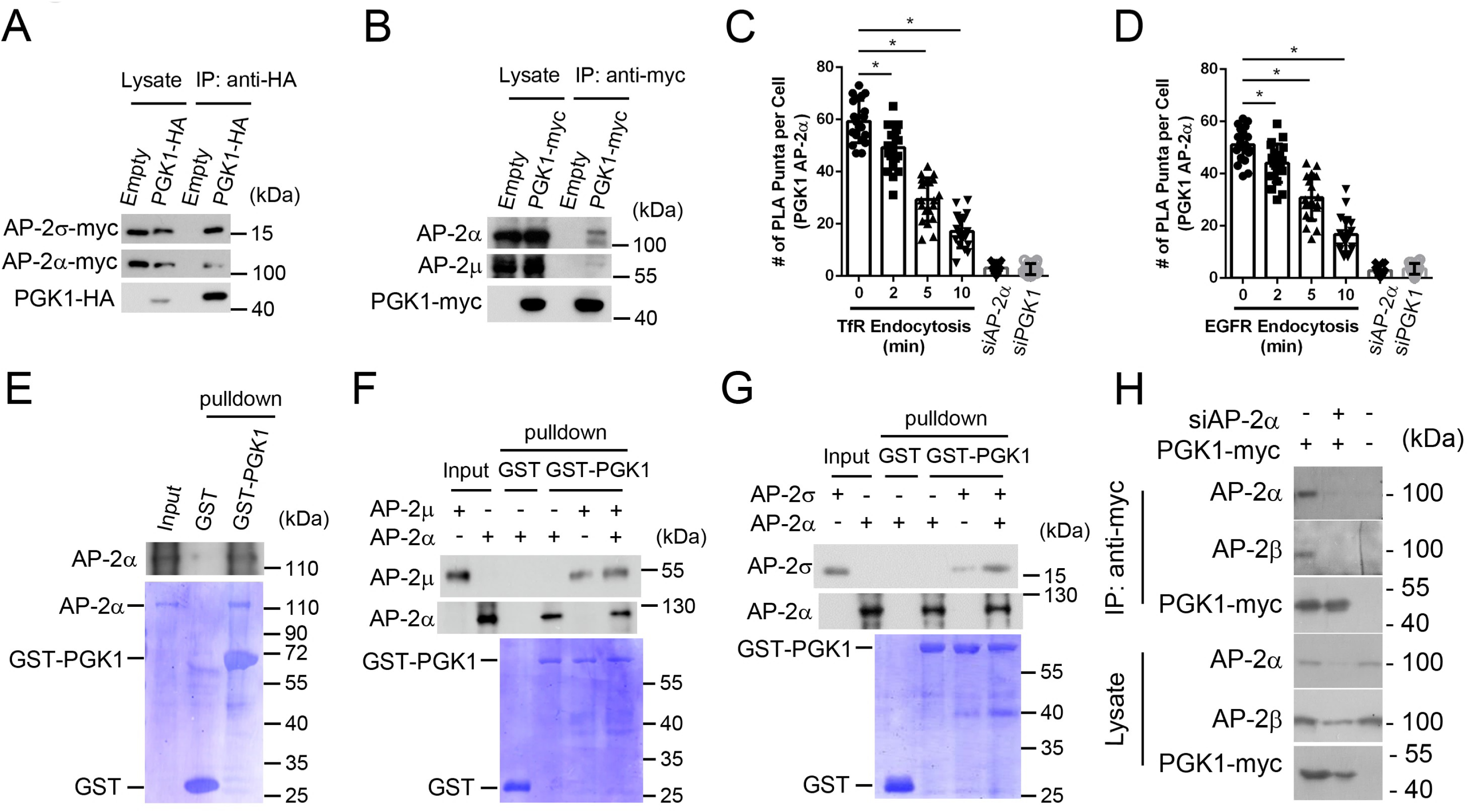
PGK1 interacts with the AP-2 complex. Data are shown as the means ± SDs. \**P*<0.05 **(A)** Coimmunoprecipitation analysis examining the associations of PGK1-HA with AP-2α-myc and AP-2σ-myc (n=3). **(B)** Coimmunoprecipitation analysis examining the association of PGK1-myc with endogenous AP-2α and AP-2μ (n=2). **(C)** PLA analysis examining the interaction between PGK1 and AP-2α during TfR endocytosis (n=20 cells, two independent experiments). One-way ANOVA with Dunnett’s test vs. 0 min: 2 min (*P*=0.0003), 5 min (*P≈*0); 10 min (*P≈*0). **(D)** PLA analysis examining the interaction between PGK1 and AP-2α during EGFR endocytosis (n=20 cells, two independent experiments). One-way ANOVA with Dunnett’s test vs. 0 min: 2 min (*P*=0.0088), 5 min (*P*=1.7×10^-13^); 10 min (*P≈*0). **(E)** Pull-down assay examining the interaction of GST-PGK1 with purified AP-2α (n=3). **(F)** Pull-down assay examining the interaction of GST-PGK1 with purified AP-2α and AP-2μ (n=3). **(G)** Pull-down assay examining the interaction of GST-PGK1 with purified AP-2α and AP-2σ (n=3). **(H)** Coimmunoprecipitation analysis examining the effect of siRNA against AP-2α on the association of PGK1-myc with endogenous AP-2β in HeLa cells (n=3).

Having demonstrated that PGK1 directly interacts with the AP-2 complex, we next analysed the consequences of altering PGK1 gene expression on the protein levels and functions of the AP- 2 complex. We first found that the expression of the AP-2α, AP-2β and AP-2μ subunits was largely diminished in siPGK1-treated cells (Fig. 3A and 3B). We also confirmed that the gene expression levels of AP-2α and AP-2μ were not significantly affected in siPGK1-treated cells (Fig. EV3A). Our findings suggest that PGK1 is required to maintain the expression of the AP-2 complex components. We next used a cycloheximide (CHX) chase assay to determine whether PGK1 knockdown influences the degradation of the endogenous AP-2 complex. Compared with that in control cells, the AP-2α protein level in PGK1-knockdown cells significantly decreased over 24 h, whereas the PGK1 protein level remained highly stable in control and siAP-2α-treated cells in the CHX chase assay (Fig. 3C and 3D). In addition, the degradation of AP-2α was blocked by treatment with the proteasome inhibitor MG132, suggesting that the lower levels of the AP-2 complex in PGK1 knockdown cells are due to proteasomal degradation (Fig. 3E and 3F). We further found that the expression of the kinase-dead mutant PGK1-T378P, like wild-type PGK1, can restore the degradation of AP-2α in siPGK1-treated cells (Fig. 3G and EV3B-3C). Collectively, these results demonstrate that PGK1 regulates the stabilization of the AP-2 complex independent of its catalytic activity.

**Figure 3.**
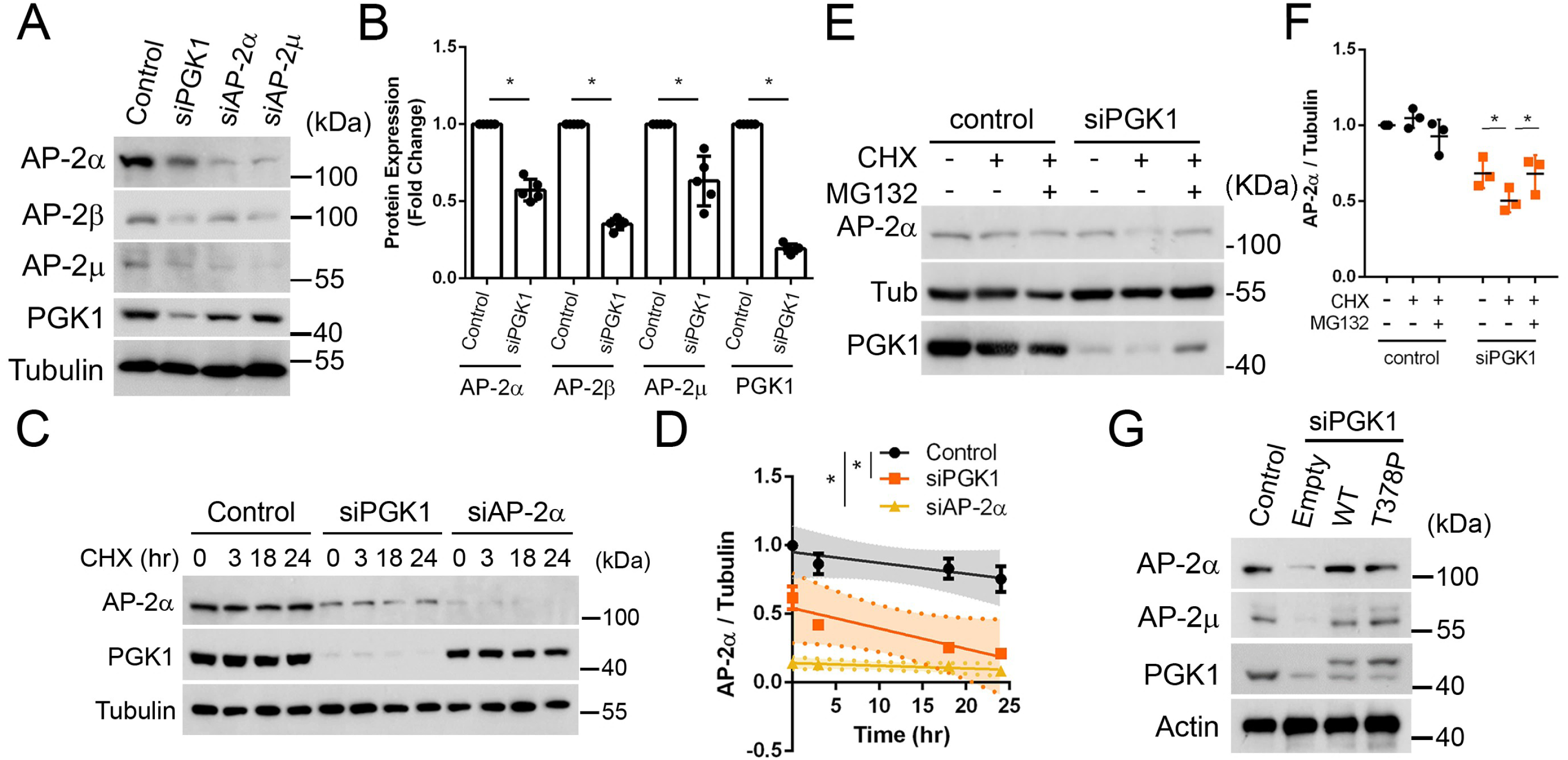
PGK1 stabilizes the AP-2 complex. Data are presented as the means ± SDs. \**P*<0.05 **(A)** Immunoblot analysis of HeLa cell lysates to examine the effects of treatment with siRNAs targeting PGK1, AP-2α, or AP-2μ. **(B)** Quantitation of the indicated protein expression levels relative to the control, normalized to tubulin levels (n=5). Unpaired two-sided Student’s t test is shown for AP-2α (*P*=8.5×10^-7^), AP- 2β (*P*=1.8×10^-10^), AP-2μ (*P*=0.00094), and PGK1 (*P*=6.8×10^-12^). **(C)** Time course analysis of AP-2α degradation upon CHX treatment for the indicated durations. **(D)** Linear regression of the AP-2α protein level. The slopes of the regression lines for control, siRNA against PGK1 and AP-2α were-0.0079,-0.0149, and-0.002, respectively (n=3). One-way ANOVA with Dunnett’s test at 24-hour time point relative to control: siPGK1 (*P*=0.0187) and siAP-2α (*P*=0.0078). **(E)** AP-2α degradation upon CHX treatment combined with MG132 treatment in control and siPGK1-treated cells (n=3). **(F)** Quantification of the AP-2α protein level normalized to the tubulin level (n=3). Two-way ANOVA with Dunnett’s test for siPGK1 cells: vehicle control versus CHX (*P*= 0.0197); CHX versus MG132 (*P*=0.0210). **(G)** Immunoblot analysis of HeLa cell lysates to examine the effects of siPGK1, PGK1 and PGK1^T378P^ (n=3).

### PGK1 acts as a cargo adaptor to promote AP-2-dependent endocytosis

To provide more direct evidence that PGK1 plays a role in protein endocytosis together with the AP-2 complex, we used multiple approaches. First, we assessed whether PGK1 and the AP-2 complex colocalized with protein cargoes during protein endocytosis. We found that both endogenous AP-2α and PGK1 significantly colocalized with the cargo proteins TfR and EGFR but not with CTxB (Fig. 4A), which is consistent with our data showing that CTxB uptake does not require PGK1 or the AP-2 complex during endocytosis (Fig. EV1D). We then examined whether the localization of one was affected upon silencing the other. We found that the colocalization of endogenous PGK1 and Tf was increased in cells treated with siRNA against AP-2α (Fig. EV4A). In cells treated with siPGK1, we also detected an increase in the colocalization of AP-2α with Tf (Fig. EV4B). These data suggest that increased PGK1 or AP-2α colocalization with cargo proteins upon AP-2α or PGK1 silencing may be the mechanism by which cells maintain protein endocytosis.

**Figure 4.**
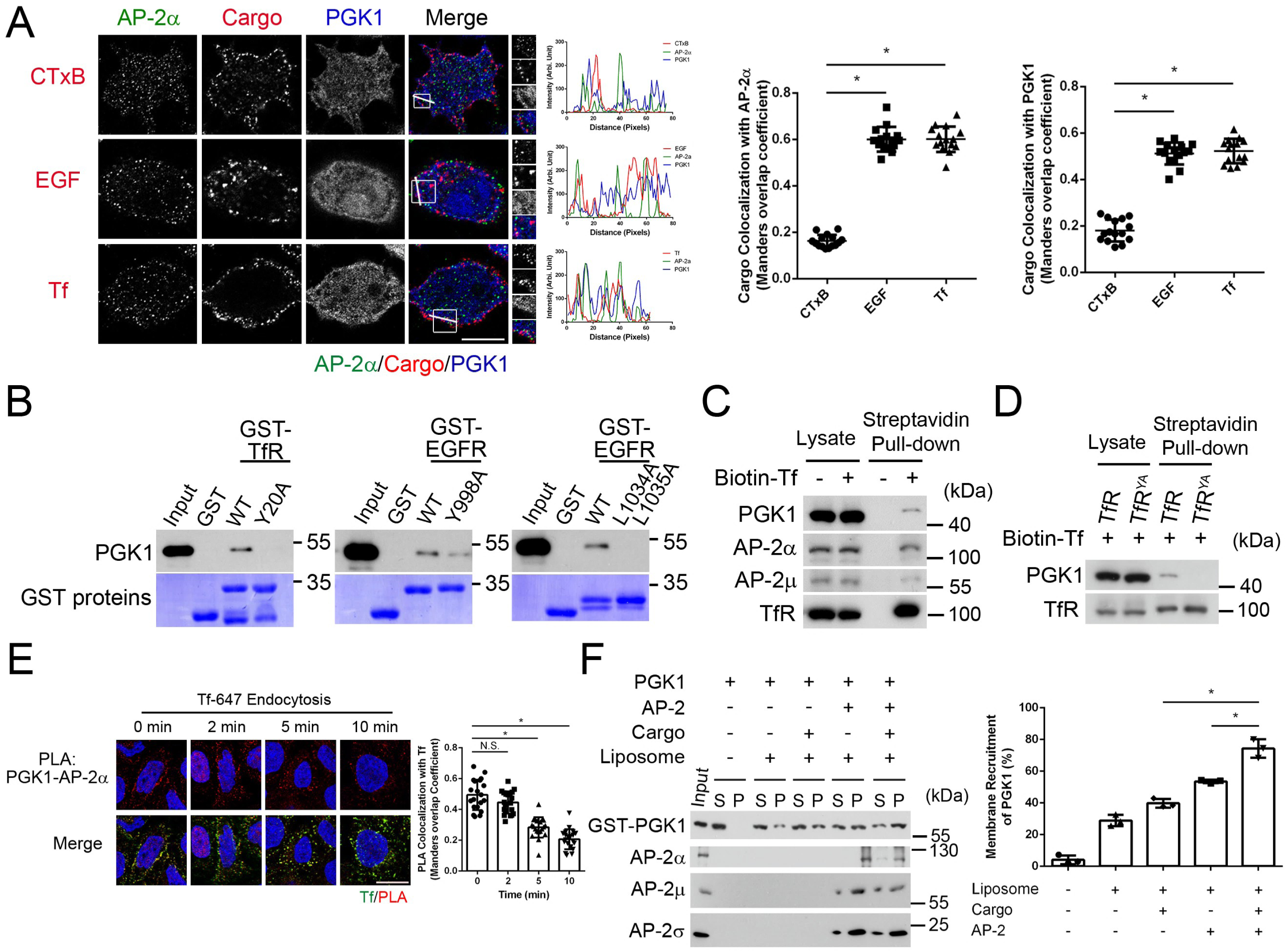
PGK1 associates with cargoes to regulate protein endocytosis. Data are shown as the means ± SDs. \**P*<0.05; NS *P*>0.05 **(A)** Immunostaining (left) of AP-2α (green), cargoes (red: CTxB, EGF, or Tf), and PGK1 (blue) at the 2-minute time point internalization; scale bar, 10 μm. Quantification (right) of colocalization between cargoes and AP-2α or between cargoes and PGK1 expressed as the Manders’ coefficient (n=15 cells, two independent experiments). One-way ANOVA with Tukey’s test relative to CTxB: AP-2α—EGF (*P*=1.1×10^-16^), Tf (*P*=6.7×10^-16^); PGK1—EGF (*P*=3.3×10^-16^), and Tf (*P*=1.1×10^-16^). **(B)** *In vitro* binding analysis of His-PGK1 and different GST-cargo fragments via GST pull-down to analyse the binding effect with different cargo fragments (n=3). **(C)** Endogenous surface TfRs associated with endogenous PGK1 and the AP-2 complex *in vivo*. Surface TfRs were bound by biotin-labeled Tf, isolated with streptavidin beads, and immunoblotted for the indicated proteins (n=3). **(D)** Myc-tagged TfR, but not TfR-YA, associated with endogenous PGK1 (n=2). **(E)** PLA analysis showing colocalization of PGK1—AP-2α PLA puncta with Tf during TfR endocytosis (n=20 cells, two independent experiments); scale bar, 10 μm. One-way ANOVA with Dunnett’s test vs. 0 min: 2 min (*P*=0.1185), 5 min (*P*=2.1×10^-13^); 10 min (*P*≈0). **(F)** Reconstitution of PGK1 recruitment to liposomes in the presence of cargo and the AP-2 complex. Representative immunoblot (left) and quantification (right) PGK1 membrane recruitment (n=3). One-way ANOVA with Dunnett’s test relative to the presence of both the cargo and the AP-2 complex: cargo only (*P*=0.0286) and AP-2 complex only (*P*=0.0295).

Second, since cargo sorting by adaptor proteins requires their direct interaction with cargo proteins, we investigated whether PGK1, like the AP-2 complex, also interacts with the cytoplasmic tails of TfR and EGFR. Previous data have shown that the dileucine motif is recognized by the AP-2σ subunit, whereas the tyrosine-based sorting signal interacts with the AP- 2μ subunit ^14,18,19^. Interestingly, in our study, PGK1 recognized both dileucine-and tyrosine-based sorting motifs, and those interactions could be specifically inhibited by mutating these sorting motifs using yeast two-hybrid assay (Fig. EV4C) and *in vitro* binding assay (Fig. 4B). Mutations of the dileucine or tyrosine-based motifs to alanine on the cytoplasmic tail of EGFR and TfR abolished binding by PGK1, indicating the specificity of the interaction (Fig. 4B). Significantly, co-incubation of PGK1 and AP-2 complex enhanced their direct interactions with cargo peptides (Fig. EV4D). We next determined whether the binding between the PGK1/AP-2 complex and surface TfRs can be detected *in vivo*. We utilized biotin-conjugated Tf bound to the cell surface and observed that the surface TfR bound to the AP-2 complex and PGK1 upon endocytosis (Fig. 4C). In addition, the transfected wild-type and kinase-dead mutant forms of PGK1 similarly associated with the surface TfR and AP-2α (Fig. EV4E), which supported our previous observation that PGK1 regulates endocytosis in a manner that is independent of its glycolytic activity. Consistently, the association between PGK1, surface EGFR and the AP-2 complex was also detected, regardless of PGK1 catalytic activity (Fig. EV4F-4G). Moreover, mutations in the tyrosine-based sorting signal of TfR significantly weakened the interaction with PGK1 (Fig. 4D).

These observations demonstrate that PGK1, like the AP-2 complex, acts as an adaptor protein to recognize sorting signals on cargo proteins to control their endocytosis.

Third, we examined the association of PGK1 and AP-2α *in situ* via PLA and detected the concurrent presence of Tf cargo protein with the location where PGK1 and AP-2α association was detected (Fig. 4E). Our data also revealed that the cooccurrence between Tf cargo protein and PGK1-AP-2α association peaked at the initial stage after the induction of endocytosis (Fig. 4E). A similar phenomenon was observed when we employed an EGF cargo protein as an alternative approach for EGFR endocytosis (Fig. EV4H). To further investigate whether cargo binding promotes the recruitment of adaptor proteins to the membrane ^20^, we prepared synthetic PI(4,5)P_2_-containing liposomes alone or liposomes with a lipid-linked tyrosine-based motif. These liposomes were then incubated with recombinant PGK1 and AP-2 components. Strikingly, PGK1 exhibited a stronger binding affinity to liposomes containing cargo peptides, and the interaction was further augmented with cargo-containing liposomes preincubated with AP-2 components (Fig. 4F), indicating that PGK1 bound to both the AP-2 complex and cargo proteins with greater affinity than PGK1 bound solely to PI(4,5)P_2_-containing liposomes. Together, our data suggest that the interaction of PGK1 with the AP-2 complex enhances cargo recognition and promotes endocytosis.

### The interaction between PGK1 and AP-2α is required for cargo sorting to support protein endocytosis

To gain insight into whether the interaction between PGK1 and AP-2α is required for protein endocytosis, we analysed the domain of AP-2α that directly interacts with PGK1. We first found that the C-terminal hinge-appendage domain was copurified with GST-PGK1 and directly interacted with PGK1, whereas the appendage domain alone significantly prevented the interaction with PGK1 (Fig. 5A and EV5A), suggesting that the hinge region of AP-2α is responsible for the interaction with PGK1. We then generated AP-2α without a hinge region (AP-2α-dH) and observed a significant reduction in PGK1 binding *in vivo* (Fig. 5B). We also confirmed that, like AP-2α, AP-2α-dH is associated with other AP-2 components (Fig. EV5B), indicating that the hinge region is specifically involved in the interaction with PGK1. Moreover, CHX chase assays revealed that, compared with AP-2α, a faster decrease in AP-2α-dH was observed (Fig. EV5C), indicating that the PGK1 binding stabilizes AP-2α. We next determined the association of surface TfR with the AP-2 complex and PGK1 upon endocytosis and found that the expression of AP-2α-dH reduced the association of both the AP-2 complex and PGK1 with TfR (Fig. 5C). PLA further confirmed these effects of AP-2α-dH, in which AP-2α-dH decreased the interaction with PGK1 *in vivo* (Fig. 5D). The colocalization of AP-2α-dH and PGK1 was also decreased upon TfR and EGFR endocytosis (Fig. EV5D-5E). Consistently, AP-2α-dH significantly inhibited both TfR and EGFR endocytosis, as indicated by a significant decrease in their internalization and colocalization with early endosomes (Fig. 5E, 5F, and EV5F-5I). Thus, these collective results revealed that the interaction of PGK1 with AP-2α through its hinge region is required for cargo sorting to promote protein endocytosis.

**Figure 5.**
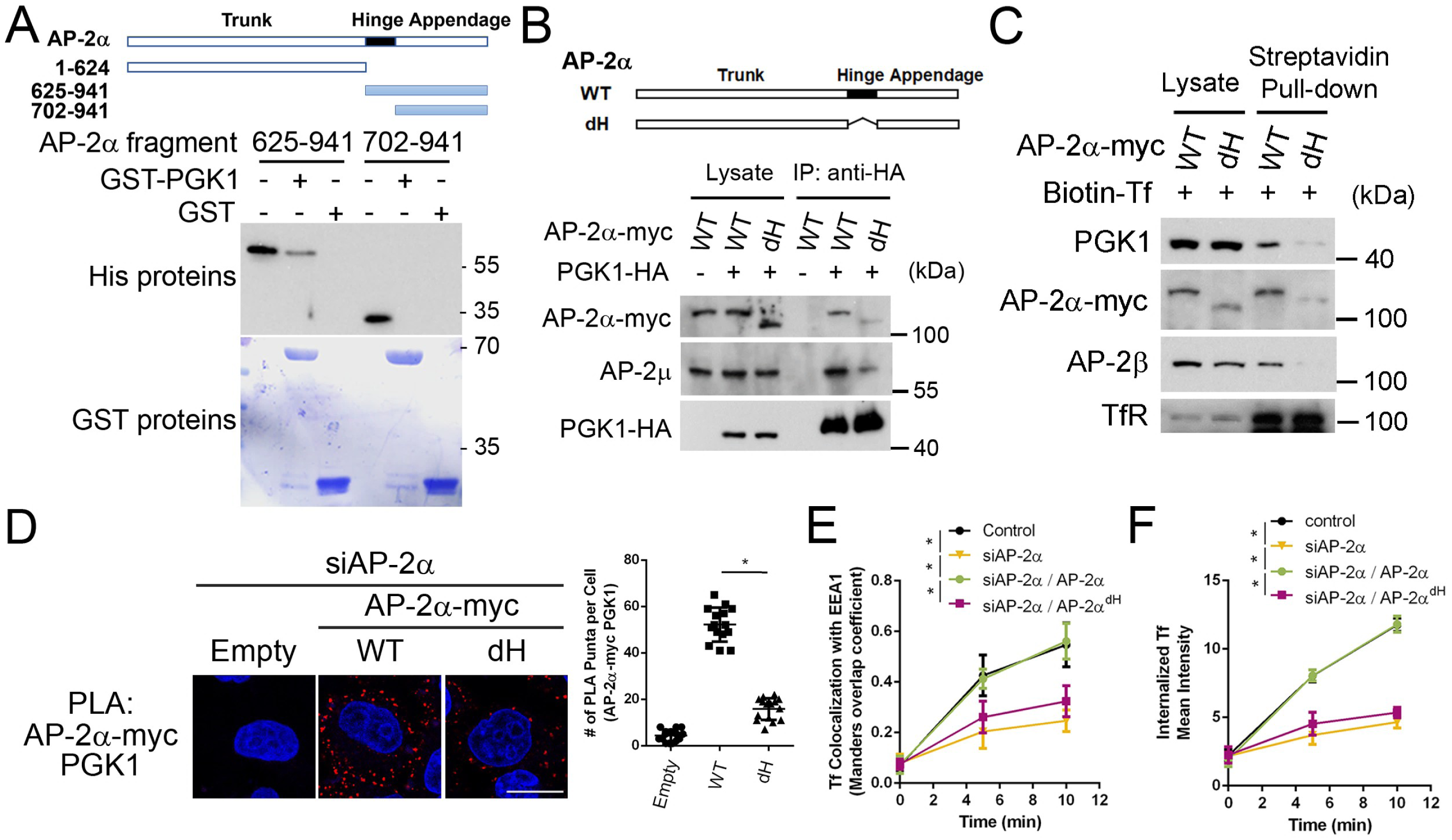
**The hinge region of AP-2**α **is required for its interaction with PGK1 and the promotion of protein endocytosis.** Data are shown as the means ± SDs. \**P*<0.05 **(A)** *In vitro* binding analysis of GST-PGK1 and different AP-2α fragments via GST pull-down to analyse the binding effect with different AP-2α fragments (n=3). **(B)** Immunoprecipitation of HA-tagged PGK1 for association with AP-2α or AP-2α^dH^ (n=3). **(C)** Surface TfRs associated with endogenous PGK1 and myc-tagged AP-2α or AP-2α^dH^ in siAP-2α-treated cells (n=3). **(D)** PLA analysis comparing the interaction of PGK1 with myc-tagged AP-2α or AP-2α^dH^ in siAP-2α-treated cells (n=15 cells, two independent experiments); scale bar, 10 μm. Unpaired two-sided Student’s t test: WT vs. dH (*P*=9.0×10^-16^). **(E)** Colocalization of Tf with EEA1, expressed as Manders’ coefficient, comparing the effects of AP-2α with AP-2α^dH^ in siAP-2α cells at 0, 5, and 10 minutes (n=15 cells, two independent experiments). One-way ANOVA with Tukey’s test at 5-minute time point: control vs. siAP-2α (*P*=9.3×10^-12^); siAP-2α vs. AP-2α (*P*=2.3×10^-11^); AP-2α vs. AP-2α^dH^ (*P*=1.6×10^-7^). **(F)** Mean intensity of internalized Tf comparing the effects of AP-2α with AP-2α^dH^ in siAP-2α cells at 0, 5, and 10 minutes (n=15 cells, two independent experiments). One-way ANOVA with Tukey’s test at 5-minute time point: control vs. siAP-2α (*P*=7.3×10^-12^); siAP-2α vs. AP-2α (*P*=7.3×10^-12^); AP-2α vs. AP-2α^dH^ (*P*=7.3×10^-12^).

We next sought to identify the specific region on PGK1 involved in the AP-2α interaction. Previous data suggested that several endocytic accessory proteins are recruited by interactions with AP-2α via the DxF motif ^21^. Notably, PGK1 contains a ^290^DKF^292^ motif that conforms to these consensus sequences. We then found that mutations in the DKF motif of PGK1 (PGK1^290–292A^) prevented its interaction with AP-2α in cells (Fig. 6A). We further found that the expression of PGK1^290–292A^ failed to reverse the degradation of AP-2α in siPGK1-treated cells (Fig. EV6A).

**Figure 6.**
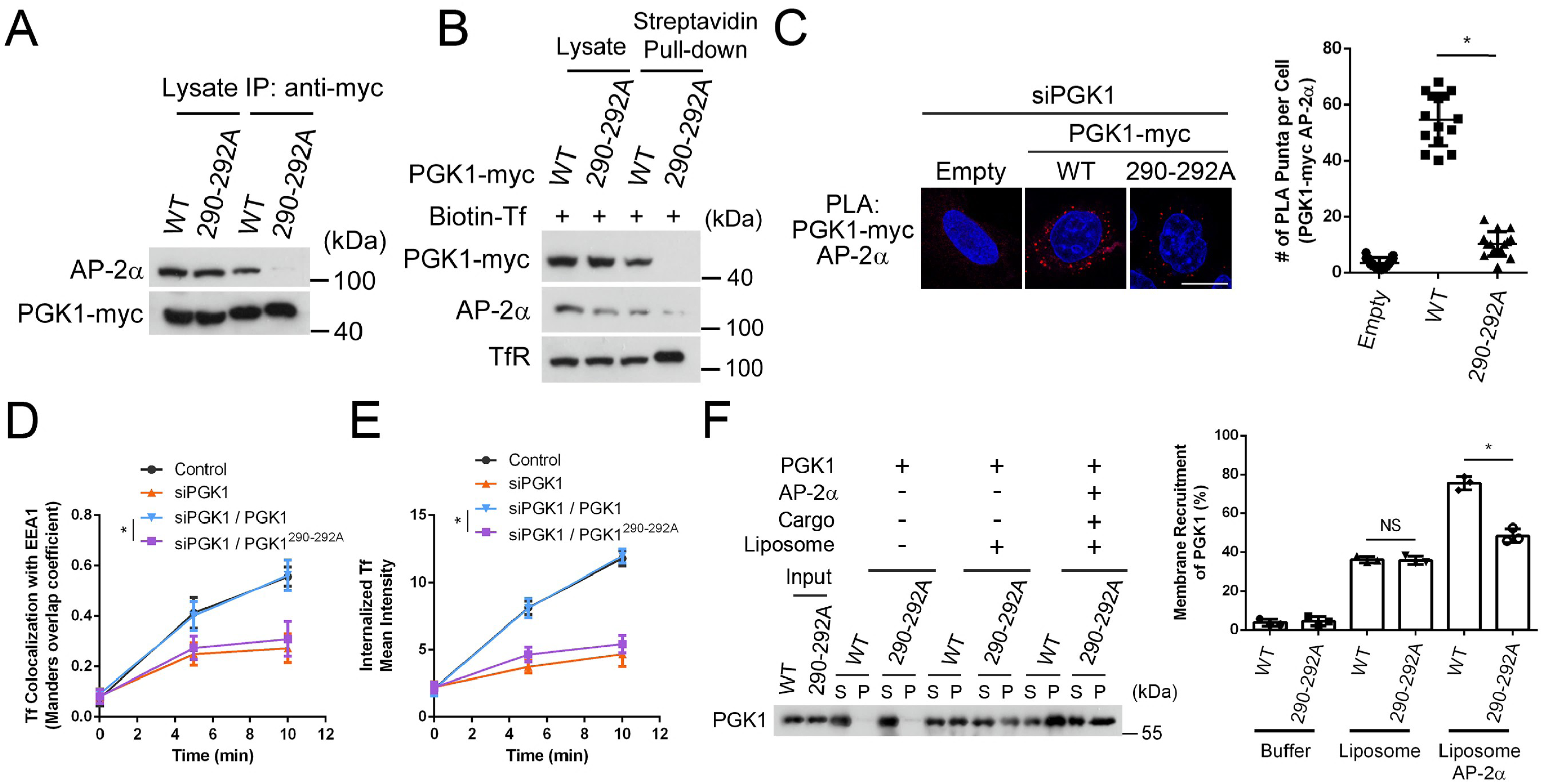
**The ^290^DKF^292^ motif of PGK1 is required for its interaction with AP-2**α **and the promotion of protein endocytosis.** Data are shown as the means ± SDs. \**P*<0.05; NS *P*>0.05 **(A)** Immunoprecipitation of myc-tagged PGK1 comparing the associations of AP-2α with PGK1 and PGK1^290-292A^ (n=3). **(B)** Endogenous surface TfRs associated with endogenous AP-2α and myc-tagged PGK1 or PGK1^290-292A^ in siPGK1-treated cells (n=3). **(C)** PLA analysis comparing the interaction of AP-2α with myc-tagged PGK1 and PGK1^290-292A^ in siPGK1-treated cells (n=15 cells, two independent experiments); scale bar, 10 μm. Unpaired two-sided Student’s t test: WT vs. 290-292A (*P*=5.3×10^-16^). **(D)** Colocalization of Tf with EEA1, expressed as Manders’ coefficient, comparing the effects of PGK1 and PGK1^290-292A^ in siPGK1 cells at 0, 5, and 10 minutes (n = 15 cells, two independent experiments). Unpaired two-sided Student’s t test: WT vs. 290-292A (*P*=3.5×10^-7^). **(E)** Mean intensity of internalized Tf comparing PGK1 and PGK1^290-292A^ in siPGK1 cells at 0, 5, and 10 minutes (n=15 cells, two independent experiments). Unpaired two-sided Student’s t test: WT vs. 290-292A (*P*=9.5×10^-15^). **(F)** Reconstitution of PGK1 recruitment to liposomes comparing the effects of PGK1 or PGK1^290-292A^. Representative immunoblott (left) and quantification (right) PGK1 membrane recruitment (n=3). Unpaired Student’s t test for WT vs. 290-292A: liposome group (*P*=0.85335); liposome in the presence of cargo and AP-2α groups (*P*=0.000691).

However, we further found that, compared with wild-type PGK1, PGK1^290–292A^ had a weaker association with cargo proteins *in vivo* (Fig. 6B and EV6B), suggesting that the interaction of PGK1 with AP-2α also promotes its association with cargo proteins. In addition, PLA further confirmed that PGK1^290–292A^ decreased the interaction with AP-2α *in vivo* (Fig. 6C). Furthermore, the expression of PGK1^290–292A^ did not restore TfR endocytosis in siPGK1-treated cells (Fig. 6D, 6E, and EV6C). Moreover, expression of PGK1^290–292A^ failed to rescue the endocytosis defect of EGFR internalization in siPGK1-treated cells (Fig. EV6D-6F). Notably, compared with wild-type PGK1, PGK^290–292A^ had a lower binding affinity for cargo-containing liposomes preincubated with AP-2 components (Fig. 6F). Thus, our data suggest that interaction between PGK1 needs to interact with AP-2α to promote protein endocytosis.

### PGK1 is part of the AP-2-clathrin coat complex

As further evidence in favour of the role of PGK1 working together with AP-2 complex to promote endocytosis, we examined whether the clathrin coat complex contains PGK1 as a coat component. We initially confirmed that AP-2α associated with PGK1 and clathrin in cells (Fig. 7A), and the interaction of PGK1 with AP-2α and clathrin was also detected via a coprecipitation experiment (Fig. 7B). We next used super-resolution confocal microscopy to observe the intracellular vesicular structure and found that a significant portion of PGK1 and AP-2α colocalized with the cargo proteins Tf and EGF (Fig. 7C and EV7A). In addition, we observed the structure of clathrin with PGK1 and the cargo proteins Tf and EGF (Fig. 7D and EV7B). To avoid potential interference from the cytosolic free portion of PGK1 during colocalization analysis, we used a pre-lysis method to eliminate glycolytic PGK1, which is abundant in the cytosol ^22^. The cells were permeabilized before fixation to sustain only the proteins associated with membranous organelles. We first found that wild-type PGK1 colocalized with clathrin, whereas the AP-2α-interacting defect mutant PGK1^290–292A^ significantly decreased the degree of colocalization (Fig. 7E). Additionally, we observed the colocalization of wild-type AP-2α with clathrin, whereas the binding of the PGK1-interacting defect mutant AP-2α-dH to clathrin was noticeably reduced (Fig. 7F). By combining super-resolution confocal microscopy with the pre-lysis method, we detected robust colocalization of the cargo proteins Tf and EGF with the PGK1-AP-2α complex in cells (Fig. EV7C-7D). We further observed the concurrent presence of clathrin with the location where PGK1 and AP-2α association was detected by PLA signals (Fig. EV7E). Moreover, compared with the decrease in TfR endocytosis in siclathrin-treated cells, TfR endocytosis was not further decreased in clathrin and PGK1 co-knockdown cells (Fig. 7G), suggesting that although the PGK1 and AP-2 complexes form co-adaptors for cargo sorting, clathrin triskelia act as the common outer coat for PGK1-mediated endocytosis. Together, these data suggest the presence of a PGK1-containing AP-2-clathrin coat complex for protein endocytosis.

**Figure 7.**
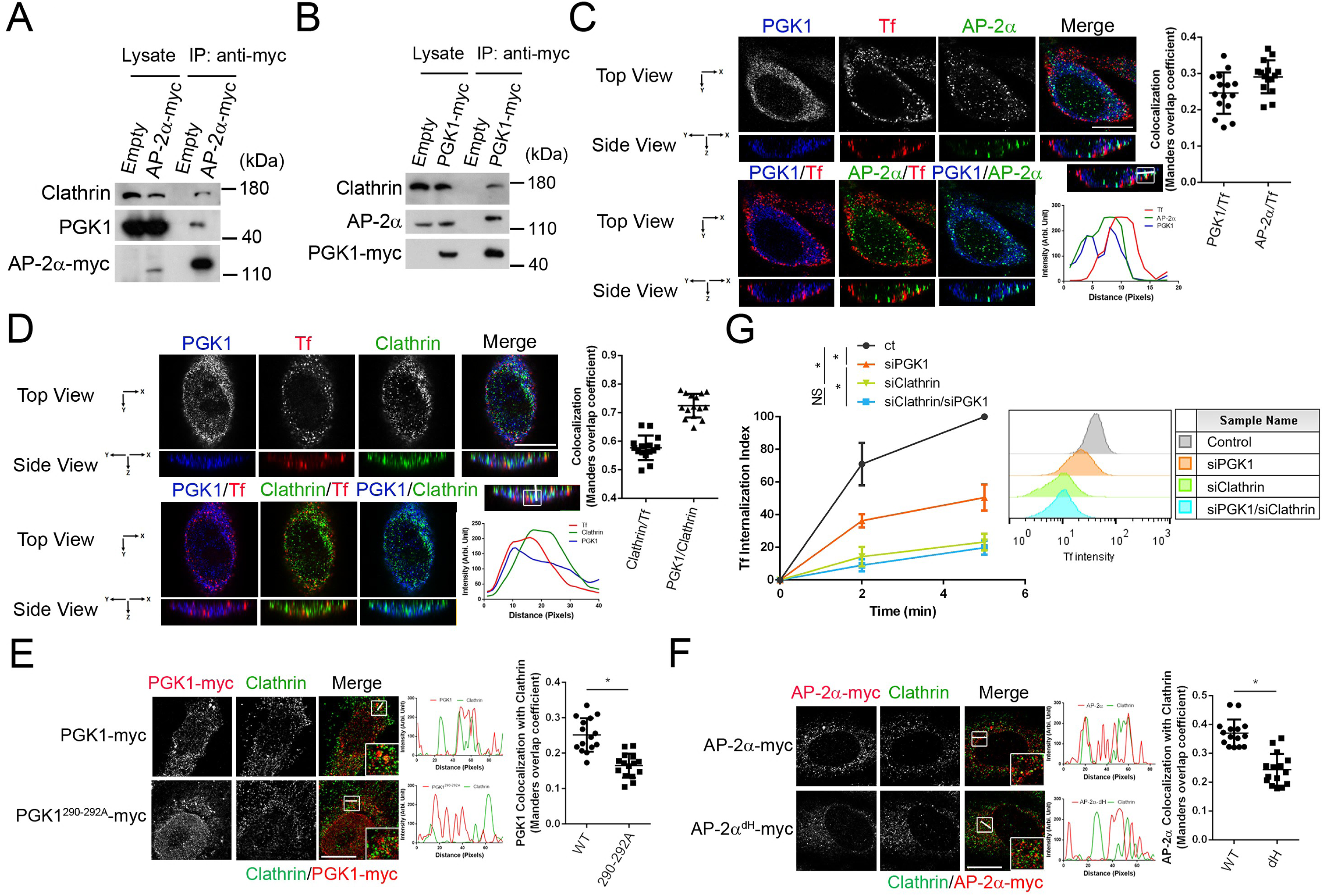
PGK1 is a part of the clathrin-coated complex during protein endocytosis. Data are shown as the means ± SDs. \**P*<0.05; NS *P*>0.05 **(A)** Coimmunoprecipitation analysis examining the association of AP-2α-myc with endogenous PGK1 and clathrin (n=3). **(B)** Coimmunoprecipitation analysis examining the association of PGK1-myc with endogenous AP-2α and clathrin (n=3). **(C)** Super-resolution confocal image showing colocalization of PGK1, AP-2α, and Tf after 2 minutes of internalization; scale bar, 10 μm. Quantification (right) of PGK1 with Tf and of AP-2α with Tf colocalization, expressed as Manders’ coefficient (n=15 cells, three independent experiments). **(D)** Super-resolution confocal image showing colocalization of PGK1, clathrin, and Tf after 2 minutes of internalization; scale bar, 10 μm. Quantification (right) of PGK1 with clathrin and clathrin with Tf colocalization, expressed as Manders’ coefficient (n=15 cells, three independent experiments). **(E)** Colocalization analysis optimized by the prelysis method demonstrating the effect of clathrin and PGK1 or PGK1^290-292A^ colocalization upon 2 minutes of internalization. Image (left; scale bar, 10 μm) and quantification (right) show the PGK1-myc colocalization with clathrin, expressed as Manders’ coefficient (n=15 cells, two independent experiments). Unpaired Student’s t test, *P*=3.38×10^-6^. **(F)** Colocalization analysis optimized by the prelysis method demonstrating the effect of clathrin and AP-2α or AP-2α^dH^ colocalization upon 2 minutes of internalization. Image (left; scale bar, 10μm) and quantification (right) show the AP-2α-myc colocalization with clathrin, expressed as Manders’ coefficient (n=15 cells, two independent experiments). Unpaired Student’s t test, *P*=2.16×10^-7^. **(G)** Internalization of Tf quantified by flow cytometry at 0, 2, and 5 minutes (n > 10,000 cells, four independent experiments). One-way ANOVA with Tukey’s test at the 5-minute time point: control vs. siPGK1 (*P*=0.0034); control vs. siClathrin (*P*=0.0002); siPGK1 vs. siClathrin/siPGK1 (*P*=0.0277); siClathrin vs. siClathrin/siPGK1 (*P*=0.5985).

To directly visualize endocytic events at the plasma membrane, we employed total internal reflection fluorescence (TIRF) microscopy which selectively detects fluorescence signals in close proximity to the plasma membrane. We first observed elevated Tf and EGF signals on the plasma membrane in cells treated with siRNA against PGK1 (Fig. 8A-8D). In siPGK1-treated cells, expression of the PGK1^290–292A^ mutant, but not wild-type PGK1, significantly failed to restore Tf and EGF internalization (Fig. 8A-8D). Consistently, we found a significant abundance of PGK1, comparable to AP-2α, in Tf-positive structures using TIRF (Fig. EV8A-8B), whereas this enrichment was absent in cells expressing PGK1^290-292A^ mutant (Fig. 8E and 8F). Together, these results indicate that PGK1 functions as a co-adaptor that cooperates with AP-2 complex to ensure proper cargo recognition and facilitate protein endocytosis.

**Figure 8.**
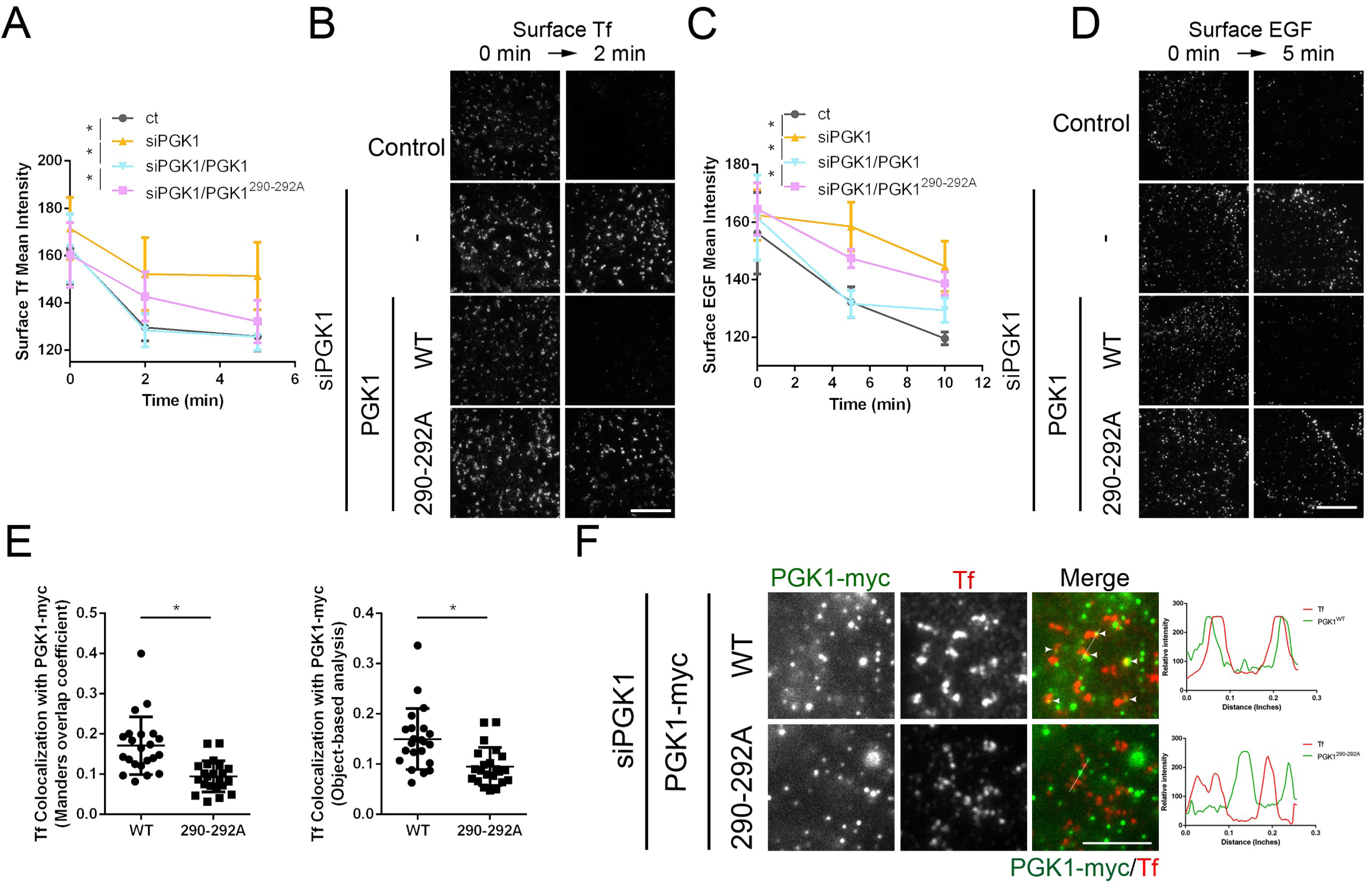
TIRF imaging demonstrated that PGK1 acts as a co-adaptor to recognize cargoes at the plasma membrane. Data are shown as the means ± SDs. \**P*<0.05 **(A)** Mean intensity of surface Tf comparing the effects of PGK1 and PGK1^290-292A^ in siPGK1-cells at 0, 2, and 5 minutes (n=20 cells, two independent experiments). One-way ANOVA with Tukey’s test at 2-minute time point: control vs. siPGK1 (*P*=7.8×10^-9^); siPGK1 vs. PGK1 (*P*=1.3×10^-9^); PGK1 vs. PGK1^290-292A^ (*P*=0.0002). **(B)** TIRF imaging representing the surface Tf; scale bar, 5 μm. **(C)** Mean intensity of surface EGF comparing the effects of PGK1 expression with PGK1^290-292A^ expression in siPGK1-cells at 0, 5, and 10 minutes (n=20 cells, two independent experiments). One-way ANOVA with Tukey’s test at 5-minute time point: control vs. siPGK1 (*P*≈0); siPGK1 vs. PGK1 (*P*≈0); PGK1 vs. PGK1^290-292A^(*P*≈0). **(D)** TIRF imaging representing the surface EGF; scale bar, 5 μm. **(E)** Quantitative results of Tf colocalization with PGK1-myc, expressed as Manders’ coefficient (left) and as object-based analysis (right) (n = 22 cells, two independent experiments). Unpaired two-sided Student’s t test: Manders’ coefficient (*P*=6.9×10^-5^); object-based analysis (*P*=0.0009). **(F)** TIRF imaging representing the colocalization of Tf and PGK1-myc; scale bar, 5 ìm. Arrows indicate the colocalization.

## DISCUSSION

We discovered that PGK1 functions together with AP-2 complex to drive efficient protein endocytosis. Elucidating this role, we initially reported that simultaneous knockdown of PGK1 and the AP-2 complex synergistically blocks protein endocytosis. We then revealed that the catalytic activity of PGK1 is not required for its regulation of endocytosis. We also noted that PGK1 directly interacts with AP-2α, stabilizing the AP-2 complex. Remarkably, unlike the AP-2 complex, in which AP-2σ and AP-2µ recognize the dileucine motif and the tyrosine-based motif, respectively, PGK1 itself can directly bind to both to promote endocytosis. The association of PGK1 and the AP-2 complex also facilitates the cargo sorting process. These observations suggest that PGK1 acts simultaneously as an interacting partner with AP-2α for the stabilization and a cargo adaptor for cargo proteins to support AP-2-dependent endocytosis. Notably, PGK1 is a key auxiliary coat component and part of the clathrin-coated complex that mediates protein endocytosis.

Our findings advance the understanding of AP-2-dependent endocytosis in multiple ways. First, the canonical tyrosine-based sorting motif and dileucine sorting signal are essential for internalization ^9^. The AP-2 complex is considered an endocytic cargo adaptor required for the internalization of certain cargo receptors, such as TfR and EGFR. Although these interactions are specific, the binding affinity needs to be increased to guarantee the proper cargo sorting step. In our case, we identified PGK1 as a cargo adaptor that works together with the AP-2 complex to recognize both sorting signals, which tends to increase the molecular avidity. Second, in addition to being an alternative endocytic accessory adaptor ^8,15,23^, PGK1 not only binds to lipids and clathrin but also associates with AP-2α to stabilize the AP-2 complex from being degraded. The dual functions, interactions with lipids and coat proteins and protective effects of PGK1 demonstrate the reciprocal role involved in protein endocytosis. Taken together, our findings suggest that PGK1 promotes TfR and EGFR endocytosis through the optimization of cargo recognition by the AP-2 complex.

The AP-2 complex facilitates endocytosis through two different stages. The initial step is to recognize PI(4,5)P_2_-containing lipids at the plasma membrane. The latter step involves a conformational change in AP-2 adaptors so that the cargo binding sites can be exposed for cargo sorting ^24,25^. The data collected in this study suggest that PGK1 can be recruited to PI(4,5)P_2_-containing liposomes and, notably, that both cargo peptides and the AP-2 complex further increase membrane recruitment. As such, the coincident detection of PI(4,5)P_2_ and cargo sorting motifs allows simultaneous recognition by the PGK1-AP-2 complex for the proper sorting step during endocytosis.

The endocytic pathway is a dynamic system responsible for lysosomal transport and membrane recycling after being internalized. Once a cargo protein is internalized from the plasma membrane, it is sorted in the early endosome. PGK1 can participate in two steps for endocytic transport: the initial step is endocytosis via the AP-2 complex revealed from this study, and the latter step is subsequent lysosomal transport of EGFR from our previous findings ^13^. Our model proposes an efficient sorting step for cargoes via the same cargo adaptor to control endocytic transport. Thus, determining how coat adaptors are tightly regulated and coordinated when one cargo protein is internalized and sorted to form different populations of transport vesicles can be more intriguing than appreciated at present.

## METHODS

### siRNA

The siRNA against PGK1, GCUUCUGGGAACAAGGUUA (siPGK1), the siRNA against AP2A1, AGAGCAUGUGCACGCUGGC (siAP-2α), the siRNA against AP2M1, AGUGGAUGCCUUUCGGGUC (siAP-2μ), and the siRNA against Clathrin, AAUUGGUCUUCGAAUUGGA (siClathrin), were prepared by Dharmacon.

### Plasmids

Wild-type PGK1, an inactive PGK1 mutant (T378P), and DKF290-292AAA (290-292A) were cloned and inserted into the BamHI and XhoI sites of pcDNA3.1-myc-His(-) (Invitrogen) and pEGFP-C3 (Clontech). HA-tagged PGK1 was also cloned and inserted into the BamHI and XhoI sites of pcDNA3.1-myc-His(-) with a stop codon after the HA tag, and mCherry was cloned and inserted into the Xba1 site of pcDNA3.1-myc-His(-), after which PGK1 was cloned and inserted into the EcoRI and BamHI sites with a stop codon. For mRFP-PGK1, PGK1 was cloned and inserted into the HindIII sites of pCMV6-AN-mRFP (OriGene) with a stop codon. PGK1 wild-type was cloned and inserted into the BamHI and XhoI sites of the wild-type pGEX4T-3 (Amersham) vector; PGK1 wild-type were cloned and inserted into the BamHI and XhoI sites of pJG4-5. AP-2α was cloned and inserted into the XhoI and HindIII sites of pcDNA3.1-myc-His(-), pIEx-6 (Novagen) and pEGFP-C3 (Clontech) and was subsequently cloned and inserted into the EcoRI and XhoI sites of pEG202 (NovoPro). For AP-2α-mCherry, mCherry was cloned and inserted into the HindIII site of pcDNA3.1-myc-His(-), after which AP-2α was cloned and inserted into the EcoRI and BamHI sites. AP-2α-GFP was obtained form addgene ((Plasmid #160214) and GFPspark-CTLB was obtained from Sino Biological (Cat: HG20112-ACG). AP-2α dH was cloned and inserted into the BamHI and XhoI sites for AP-2α 1-624 and the XhoI and HindIII sites for AP-2α 702-941 of pcDNA3.1-myc-His(-). AP-2μ was cloned and inserted into the XhoI and EcoRV sites of pcDNA3.1-myc-His(-) and pET32a (Invitrogen). AP-2σ was cloned and inserted into the XhoI and BamHI sites of pcDNA3.1-myc-His(-) and pET32a (Invitrogen). For truncated AP-2α: 1-624, 625-941, and 702-941 were cloned and inserted into the XhoI and BamHI sites of pET32a and pET30a (Invitrogen). Cargoes peptides used in GST-pulldown assay: For the TfR cytosolic tail, TfR1-61 (TfR cyto) wild-type and Y20A mutant strains were cloned and inserted into pGEX4T-3. For the EGFR cytoplasmic tail containing tyrosine motif, EGFR 980-1017 wild-type and Y998A mutant strains were cloned and inserted into the BamHI sites of pGEX4T-3. For the EGFR cytoplasmic tail containing di-leucin motif, EGFR 1013-1054 wild-type and L1034AL1035A mutant strains were cloned and inserted into the BamHI sites of pGEX4T-3.

Cargoes peptides used in Yeast-two hybrid: For the TfR cytosolic tail, TfR1-61 (TfR cyto) wild-type and Y20A mutant strains were cloned and inserted into the EcoRI and XhoI sites of pEG202. For the EGFR cytoplasmic tail containing tyrosine motif, EGFR 992-1005 wild-type, and Y998A mutant strains were cloned and inserted into the BamHI sites of pEG202. For the EGFR cytosolic tail containing di-leucin motif, EGFR 1018-1045 wild-type, and L1034AL1035A mutant strains were cloned and inserted into the BamHI sites of pEG202.

### Antibodies

The following antibodies were used: anti-EEA1 (Cell Signalling 2411, immunofluorescence, 1:100/western blotting, 1:1000), anti-Myc (Cell Signaling 2276, immunofluorescence, 1:100/western blotting, 1:1000), anti-6×His (Santa Cruz sc-803, western blotting, 1:500), anti-HA (Santa Cruz sc-7392, western blotting, 1:1000), anti-PGK1 (Santa Cruz sc-130335, immunofluorescence, 1:100/western blotting, 1:2000) (GeneTex GTX107614, immunofluorescence, 1:100), anti-AP-2α (BD Transduction Laboratories 610502, immunofluorescence, 1:100/western blotting, 1:1000), anti-AP-2μ (BD Transduction Laboratories 611351, western blot, 1:500), anti-AP-2β (Santa Cruz sc-74423, western blotting, 1:1000), anti-AP-2σ (ORIGENE TA373563, western blotting, 1:500), anti-clathrin heavy chain (Cell Signaling 2410, western blotting, 1:1000), anti-α-tubulin (Cell Signaling 3873, western blotting, 1:5000), anti-EGFR (Cell Signaling 4267, immunofluorescence, 1:100/western blotting, 1:1000), anti-TfR (Santa Cruz sc-65882, western blotting, 1:500), secondary antibodies for western blotting (Merck Millipore): peroxidase-conjugated goat anti-mouse IgG (AP124P, 1:5000) and anti-rabbit IgG (AP132P, 1:5000), secondary antibodies for immunofluorescence (Jackson ImmunoResearch): Alexa Flor® 488 (115-545-062)/594 (115-585-062)/647 (115-605-062)-conjugated AffiniPure goat anti-mouse and Alexa Flor® 488 (711-545-152)/594 (711-585-152)/647 (711-605-152)-conjugated AffiniPure donkey anti-rabbit.

### Cell culture

All mammalian cells were cultured at 37°C with 5% CO_2_. HeLa (ATCC CCL-2) and 293T (ATCC CRL-3216) cells were cultured in Dulbecco’s modified Eagle’s medium (DMEM) with high glucose (Cytiva SH30022.02) supplemented with 10% fetal bovine serum (FBS; Corning™ 35010CV) and 1% penicillin‒streptomycin solution (PS; Cytiva SV30010). Unless otherwise indicated, all experiments were performed using HeLa cells. For siRNA transfection, PepMute^™^ siRNA transfection reagent (SignalGen^®^ SL100566) was used to silence target genes. For plasmid transfection, FuGENE^®^ 6 Transfection Reagent (Promega E2691) was used for transient expression.

Insect ExpiSf9™ cells (Gibco™ A35243) were cultured at 25°C in ExpiSf™ CD medium (Gibco™ A3767801). For plasmid transfection, polyethylenimine, linear, MW 25000, transfection grade (PEI 25K™; polysciences 23966) was used for transient expression.

The yeast YEM1α (*MATα*, *trp1*, *his3*, *leu2*, *6ops*-*LEU2*, *2ops*-*LacZ*) strain was used to conduct a yeast two-hybrid assay and was maintained in yeast extract peptone dextrose (YPD) medium containing 1% yeast extract, 2% peptone, 2% glucose or synthetic dextrose (SD) medium containing 0.67% yeast nitrogen base (YNB), 2% glucose, and NaOH (pH=8) with an appropriate amino acid mixture. For plasmid transformation, a frozen-EZ yeast transformation II kit was used. To express the desired proteins, 2% galactose was used to replace glucose and induce protein expression under the gal promoter.

### Endocytosis assay

Alexa Fluor dye-conjugated cargoes, including Alexa Fluor 488/555/647-conjugated transferrin, Alexa Fluor 488/555/647-conjugated EGF, and Alexa Fluor 555-conjugated CTxB, were obtained from Invitrogen.

To examine cargo transport from the plasma membrane to the early endosome, the cells were cultured in serum-free medium for one hour and then allowed to bind to the cargo (5 μg/mL for Tf, 0.1 μg/mL for EGF, or 5 μg/mL for CTxB) for one hour at 4°C. The cells were then incubated at 37°C and examined at the indicated time points. For those needed to measure the internalized Tf or EGF intensity, cells were first washed by the acid buffer, 0.2M acetic acid and 0.5M NaCl, and then fixed. Fixed cells were stained for EEA1 and observed via confocal microscopy to assess the colocalization of the cargoes and EEA1. Images were analysed with Fiji-ImageJ2.

### Internalization index measured by flow cytometry

To examine Tf internalization, serum-starved cells were incubated with Alexa Fluor 488-conjugated Tf at 4°C for one hour. To measure Tf levels, the cells were fixed and collected in Falcon^®^ (Cat. No. 352052). To measure internalized Tf levels, cells were then incubated at 37°C.

To quantify Tf internalization at constant 37°C, serum-starved cells were incubated with Alexa Fluor 488–conjugated Tf and maintained at 37°C for 0, 2, and 5 minutes, following by acidic buffer quenching. As a negative control, cells were pre-treated with 80 µM dynasore for 10 min to inhibit endocytosis^17^.

For trypan blue quenching, fixed cells were treated with 2% trypan blue in phosphate-buffered saline (PBS) to quench surface Tf ^26^. For acidic buffer quenching, cells were washed with acidic buffer (0.2M acetic acid and 0.5M NaCl) before fixation. The fluorescence intensity of approximately 10,000 cells at 495 nm was analyzed using a BD FACSCalibur flow cytometer. Data were analysed with FlowJo version 10.

For trypan blue-treated groups, the internalization index was calculated as: *Internalization Index*=*(I_TB_(t)-I_TB_(0))/I(0)*. where *I* represents relative fluorescence intensity at 495 nm, *TB* indicates trypan blue treatment, and *t* represents incubation time (minutes).

For acidic wash groups, the internalization index was calculated as: *Internalization Index=(I_A_(t)-I(0))/I(t)*, where A indicates cells washed with acidic buffer.

### Recombinant protein purification

*Escherichia coli* (*E. coli*) strain BL21 (Novagen) was transformed with plasmids, pGEX-4T-3 for the GST-tag, and pET30a or pET32a for the His-tag. The transformed E. *coli* were harvested at the exponential growth phase and induced to express recombinant proteins with 500 μM isopropyl-β-thiogalactoside (IPTG) at 37°C for three hours. For the GST tag, GST-tagged PGK1 or GST-tagged cargoes, the lysates were purified in PBS with glutathione-Sepharose 4B (GE Healthcare 17075605), and the coexpression of GST-PGK1 and truncated AP-2α also followed the protocol of GST protein purification. For His-tagged truncated AP-2α, lysates were lysed and pulled down with nickel affinity resin (Thermo 88222) in 50 mM NaH_2_PO_4_-NaOH (pH=8.0), 300 mM NaCl, and 10 mM imidazole, followed by elution with 250 mM imidazole. For protein storage, the buffer was exchanged for PBS by dialysis (Spectra/Por 4 Dialysis Tubing, 12–14 kDa MWCO; Repligen 132700). For His-tagged AP-2μ and AP-2σ, lysates were lysed and pulled down with nickel affinity resin (Thermo 88222) in 20 mM Tris-HCl (pH=8.0), 500 mM NaCl, 5 mM imidazole, and 6 M urea, after which the proteins were eluted with 250 mM imidazole without urea.

For His-tagged full-length AP-2α, pIEx-6 was transiently expressed in ExpiSf9™ cells (Gibco™ A35243). Lysates were lysed and pulled down with nickel affinity resin in 50 mM NaH_2_PO_4_-NaOH (pH=8.0) with 300 mM NaCl and 10 mM imidazole, followed by elution with 250 mM imidazole. For protein storage, the buffer was exchanged with PBS by dialysis.

### Pull-down assay

To assess the binding of PGK1 to the AP-2 components, GST-PGK1 with glutathione beads was incubated with the AP-2 components in 50 mM KCl, 25 mM HEPES-KOH (pH=7.2) with 2.5 mM Mg(OAc)_2,_ and a protease inhibitor for an hour at 4°C. The beads were washed following analysis via gel electrophoresis, followed by either western blotting with an anti-His antibody or Coomassie blue staining.

To assess the binding of PGK1 to truncated AP-2α, GST-PGK1 with glutathione beads was incubated with truncated AP-2α in 50 mM HEPES-KOH (pH=7.2) with 300 mM NaCl, 1 mM EDTA, 0.5% NP40 and protease inhibitor for an hour at 4°C. The beads were washed following analysis via gel electrophoresis, followed by either western blotting with an anti-His antibody or Coomassie blue staining.

### Mass spectrometry analysis

Proteins isolated from the immunoprecipitation assay were separated by 12.5% SDS-PAGE and visualized using Coomassie Brilliant Blue G-250 staining. The gel bands were excised and in-gel tryptic digestion was performed according to previously established protocols ^27^. The resulting peptide mixtures were extracted with acetonitrile to achieve a final concentration of 50% and subsequently dried using a SpeedVac. For the liquid chromatography tandem-mass spectrometry analysis, each peptide mixture was reconstituted in HPLC buffer A (0.1% formic acid, Sigma) and introduced into a trap column (Zorbax 300 SB-C18, 0.3 × 5 mm), where they were washed with buffer A. The desalted peptides were then separated on a 10 cm analytical C18 column with an inner diameter of 75 μm. The liquid chromatography system was linked to a linear ion trap mass spectrometer (Orbitrap Elite, Thermo Fisher Scientific). Full-scan mass spectrometry was conducted with the Orbitrap over a mass range of 400–2000 Da, achieving the detection of intact peptides at a resolution of 30,000. To identify the proteins, the raw data from the mass spectrometry were processed using Proteome Discoverer software (version 1.4, Thermo Fisher Scientific). The obtained mass spectra were searched against the SwissProt database (Taxonomy: Homo sapiens) through the Mascot search engine (Matrix Science). The search parameters included trypsin as the digestion enzyme, allowing up to two missed cleavages. Carbamidomethylation (C) was set as a static modification, while dynamic modifications included oxidation (M), N-acetylation (protein), and Gln to pyro-Glu (N-term Q). The tolerances were set at 10 ppm for MS and 0.7 Da for MS/MS.

### Immunofluorescence staining and PLA

Cells seeded on coverslips were fixed with 2% paraformaldehyde in PBS for 10 minutes and permeabilized in PBS with 3% BSA and 0.1% saponin for 20 minutes at room temperature. Primary and secondary antibodies were diluted in PBS with 10% FBS, 0.02% NaN_3_, and 0.2% saponin and incubated with coverslips for an hour at room temperature. After immunostaining, the coverslips were mounted on slides with Fluoromount-G™ Mounting Medium with DAPI (Invitrogen™ 00-4959-52). To prelyse the cell membrane, release cytosolic PGK1, and increase the endosomal PGK1 signal, coverslips were pretreated with 0.0625% saponin in PBS for 2 minutes at room temperature before fixation ^22^.

For the PLA, following the protocol of the immunostaining and PLA kit (Duolink® *In Situ*, Sigma), cells seeded on coverslips were fixed with 2% paraformaldehyde in PBS for 10 minutes and permeabilized in PBS with 3% BSA and 0.1% saponin for 20 minutes at room temperature. Primary antibodies were diluted in PBS with 10% FBS, 0.02% NaN_3_, and 0.2% saponin for an hour at room temperature and washed twice with wash buffer A. The cells were incubated with a mixture of Duolink® *in situ* PLA® probe anti-rabbit PLUS (Duolink® DUO92002) and Duolink® *in situ* PLA® probe anti-mouse MINUS (Duolink® DUO92004) for an hour at 37°C and washed twice with wash buffer A. To ligate the two probes, the cells were incubated with ligation buffer and ligase for 30 minutes at 37°C and washed twice with wash buffer A. To amplify the PLA signal, the cells were incubated with amplification buffer and polymerase for 100 minutes at 37°C and washed twice with wash buffer B. The cells were then mounted on slides with Fluoromount-G™ Mounting Medium with DAPI (Invitrogen™ 00-4959-52).

### Yeast two-hybrid assay

Yeast cells transformed with pJG4-5 and pEG202 were plated on a synthetic galactose (SG) plate (2% galactose) with His, Trp dropout (-His-Trp), and His, Trp, and Leu dropout (-His-Trp-Leu) plates were incubated at 30°C for the appropriate time. Protein‒protein interactions enabled the growth of yeast cells on His, Trp, and Leu dropout plates.

### Immunoprecipitation

Mammalian cells were lysed in 50 mM Tris-HCl (pH=7.4), 150 mM NaCl, 0.5% Triton X-100, 20 mM NaF, protease inhibitors, and phosphatase inhibitors. The supernatants were incubated with anti-c-myc agarose conjugate (Sigma A7470) or anti-HA agarose conjugate (Sigma A2095) at 4°C for two hours. The beads were washed three times and subjected to western blot analysis or mass spectrometry.

### Cycloheximide chase assay

The cells were treated with 10 μg/mL CHX (Sigma C1988) for the indicated time and then were lysed in 50 mM Tris-HCl (pH=7.4), 150 mM NaCl, 5 mM EDTA, 1% Triton X-100, protease inhibitors, and phosphatase inhibitors, followed by western blot analysis. To determine whether degradation occurs through the proteasome, the cells were coincubated with 30 mM MG132 (Cell Signaling, 2194) and 10 μg/mL CHX for 18 hours, lysed, and then subjected to analysis via western blotting.

### Quantitative PCR (qPCR)

Total RNA was isolated and transformed into cDNA with a reverse transcription kit (PROtech PT-RT-KIT) and Oligo (dT)_18_ primers. Gene expression was quantified via Bio-Rad CFX Connect qPCR using Fast SYBR™ Green Master Mix (Thermo 4385612) with the indicated gene primers. The gene expression of ACTIN was used as an internal control. The primers for ACTIN were forwards-TGATCTTCATTGTGCTGGGTG and reverse-ACATCCGCAAAGACCTGTAC. The primers for AP2A1 were forwards, CGGAACTGTAAGAGCAAAGAGG, and reverse, CCAAGCAGGAAGATGAAAAGC. The primers for AP2M1 were forwards CAAGCAAGAGCGGGAAGC and reverse CCATCTGGCGGGATAAAGC.

### Streptavidin pull-down assay

Serum-starved cells were incubated with 5 μg/mL biotin-labelled transferrin and 100 ng/mL biotin-labelled EGF at 4°C for one hour and then washed with unbound. The cells were lysed in 50 mM Tris-HCl (pH=7.4), 150 mM NaCl, 0.5% Triton X-100, 20 mM NaF, protease inhibitors, and phosphatase inhibitors. The supernatants were incubated with streptavidin-agarose (Merck Millipore 16126) at 4°C for one hour. The beads were washed three times and subjected to western blot analysis.

### Liposome recruitment assay

The liposomes contained 50% phosphatidylcholine (DOPC; Avanti® 850375C), 40% phosphoethanolamine (DOPE; Avanti® 850725C), and 10% 18:1 phosphatidylinositol-4,5-bisphosphate (PI(4,5)P_2_; Avanti® 850155P). To capture the peptides, the liposomes contained 40% DOPC, 40% DOPE, 10% PI(4,5)P_2_, and 10% 1,2-dioleoyl-sn-glycero-3-[(N-(5-amino-1- carboxypentyl) iminodiacetic acid)succinyl] (nickel salt) (DGS-NTA(Ni); Avanti® 790404C) ^20^. Liposomes were rehydrated in 25 mM HEPES-KOH (pH 7.2), 50 mM KCl, 2.5 mM Mg(OAc)_2_, and 200 mM sucrose overnight and passed through a 400 nm filter with a miniextruder. His-diLeu peptides (HHHHHHVEPLTPSGEAPNQA<u>LL</u>) and His-Tyr peptides (HHHHHHSAFSNLFGGEPLS<u>Y</u>TRFSL) were synthesized by our synthesis core facility (Synthesis Core Facility, Institute of Biological Chemistry, Academia Sinica), and 20 μg liposomes were preincubated with 500 nM cargo peptides at 37°C for an hour if needed. After preincubation, 20 μg of liposomes were incubated with 300 nM GST-PGK1, 300 nM His-AP-2α, 300 nM AP-2μ, and 300 nM AP-2σ, as indicated, in 25 mM HEPES-KOH (pH 7.2), 50 mM KCl, and 2.5 mM Mg(OAc)_2_ at 37°C for an hour. After centrifugation, the liposomes were pelleted, and the supernatants were collected and precipitated with 10% trichloroacetic acid (Sigma T6399). Both the supernatants and pellets were subjected to western blot analysis.

### TIRF microscopy and image analysis

All the confocal images were obtained with a Nikon ECLIPSE Ti2 TIRF microscope with a Plan Apochromat TIRF 100X/1.49 NA objective. To examine the Tf level on the plasma membrane, serum-starved cells were incubated with Alexa Fluor 555-conjugated Tf at 4°C for one hour. To measure the initial surface Tf level, cells were fixed at 4°C. To measure the remaining level of surface Tf after internalization, the cells were shifted to 37°C and then fixed.

Cells were observed in PBS via TIRF microscopy. For colocalization quantification, images were analysed via *Fiji-ImageJ2*. *Apply_DOG_Filtering.py* was used to subtract the background fluorescence, followed by threshold adjustment using *threshold_adjuster*. Colocalization for Manders’ coefficient and object-based analysis were performed using *JACoP v2.0* ^28^.

### Confocal microscopy and image analysis

All the confocal images were obtained with a ZEISS LSM 900 confocal microscope with a Plan Apochromat 63X/1.4 NA objective, and super-resolution confocal images were obtained with a ZEISS LSM 900 with Airyscan 2. Images were analysed via Fiji-ImageJ2. Colocalization was measured via Coloc2 (<u>imagej.net/plugins/coloc-2</u>), expressed as the Manders’ overlap coefficient, and the intensity profile was measured via an RGB profiler (<u>imagej.net/plugins/rgb-profiler</u>).

### Statistical analysis

The quantitative data are shown as the means +/- SEMs. Statistical significance was determined using Excel, Prism software, and R-4.4.1.

## ACKNOWLEDGEMENTS

We thank the Synthesis Core Facility, Institute of Biological Chemistry, Academia Sinica, for the peptide synthesis and the Imaging Core at the First Core Labs, National Taiwan University College of Medicine, for image acquisition using the Zeiss Elyra7 TIRF microscope and tracking analysis using Imaris. We acknowledge the technical assistance of the IBS facility at the Institute of Biological Chemistry, Academia Sinica, Taiwan. This work is supported by grants from the National Science and Technology Council (NSTC), Taiwan (111-2636-B-002-029, 112-2636-B- 002-006, 113-2636-B-002-002, and 114-2311-B-002-023) to J.-W. H., the Yushan Fellow Program by the Ministry of Education (MOE), Taiwan (MOE-109-YSFAG-0003-002-P1) to J.-W. H.

## AUTHOR CONTRIBUTIONS

SLC performed the majority of experiments including transport assay, colocalization analysis, protein-protein interaction assay, and TIRF studies. JWH performed transport assay and PLA assay. YTC conducted yeast two-hybrid analysis, protein-protein interaction analysis, and super-resolution confocal analysis. CYH supported the yeast two-hybrid assay. CJY conducted and annotated the mass spectrometry analysis. JWH and SLC designed the study and analysed the data. JWH supervised the study and experiments. JWH and SLC wrote the manuscript with input from all other authors.

## EXPANDED VIEW FIGURE LEGENDS

**Figure EV1.**
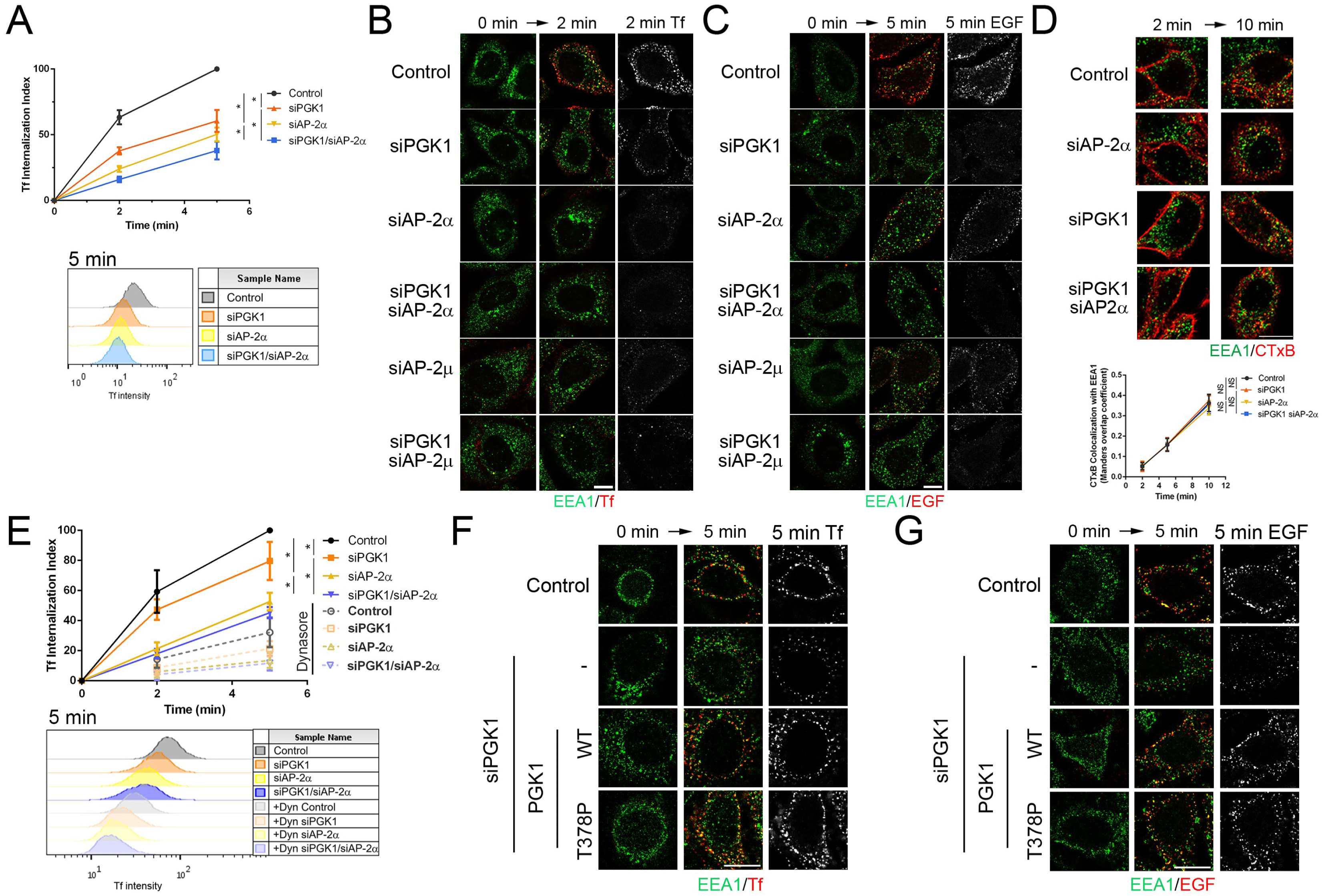
PGK1 regulates AP-2-mediated endocytosis. Quantitative analyses of cells treated with the indicated siRNAs are shown as the means ± SDs. \**P*<0.05; NS *P*>0.05 **(A)** Internalization of Tf quantified by flow cytometry at 0, 2, and 5 minutes (n > 10,000 cells, four independent experiments). One-way ANOVA with Tukey’s test at the 5-minute time point: control vs. siPGK1 (*P*=8.7×10^-7^); control vs. siAP-2α (*P*=7.0×10^-9^); siAP-2α vs. siPGK1/siAP- 2α (*P*=0.025); siPGK1 vs. siPGK1/siAP-2α (*P*=4.8×10^-6^). **(B)** Representative image of internalized Tf (red) colocalized with EEA1 (green); scale bar, 10 μm. **(C)** Representative image of internalized EGF (red) colocalized with EEA1 (green); scale bar, 10μm. **(D)** Image (left; scale bar, 10 μm) and quantification showing CTxB (red) colocalized with EEA1 (green), expressed as Manders’ coefficient (n=15 cells, two independent experiments). One-way ANOVA with Tukey’s test for the 10-minute time point: control vs. siPGK1 (*P*=0.7771); control vs. siAP-2α (*P*=0.3296); siPGK1 vs. siPGK1/siAP-2α (*P*=0.4596); siAP-2α vs. siPGK1/siAP- 2α (*P*=0.6389). **(E)** Tf Internalization at constant 37°C quantified by flow cytometry (n>10,000 cells, four independent experiments); dynasore treatment as negative control. One-way ANOVA with Tukey’s test at the 5-minute time point: control vs. siPGK1 (*P*=0.022); control vs. siAP-2α (*P*=1.7×10^-9^); siAP-2α vs. siPGK1/siAP-2α (*P*=0.0043); siPGK1 vs. siPGK1/siAP-2α (*P*=0.0106). **(F)** Representative image of internalized Tf (red) colocalized with EEA1 (green); scale bar, 10 μm. **(G)** Representative image of internalized EGF (red) colocalized with EEA1 (green); scale bar, 10μm.

**Figure EV2.**
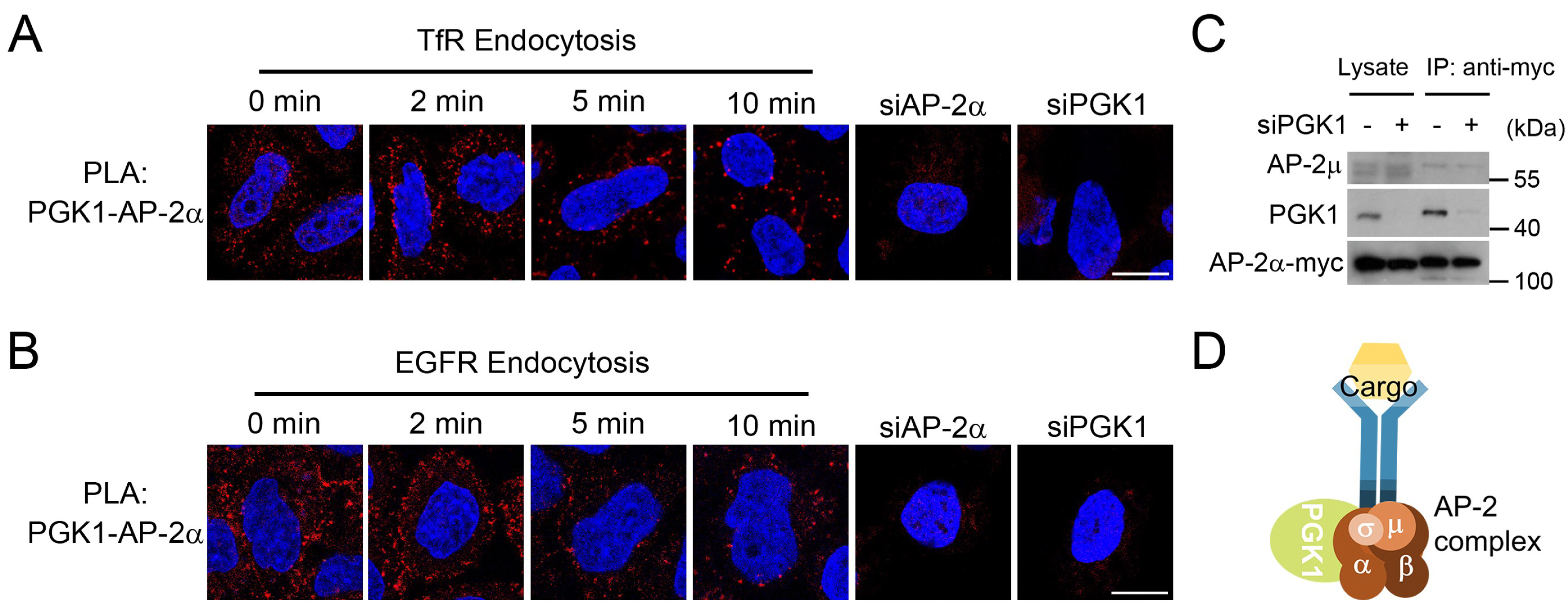
PGK1 interacts with AP-2α **(A)** PLA analysis showing the interaction between PGK1 and AP-2α during TfR endocytosis. Red indicates PLA puncta and the blue indicates nuclear staining; scale bar, 10 μm. **(B)** PLA analysis showing the interaction between PGK1 and AP-2α during EGFR endocytosis. Red indicates PLA puncta and the blue indicates nuclear staining; scale bar, 10 μm. **(C)** Co-immunoprecipitation analysis examining the effect of PGK1 siRNA on the association of myc-tagged AP-2α with endogenous AP-2μ (n=3). **(D)** A representative model for PGK1 and AP-2 complex interactions.

**Figure EV3.**
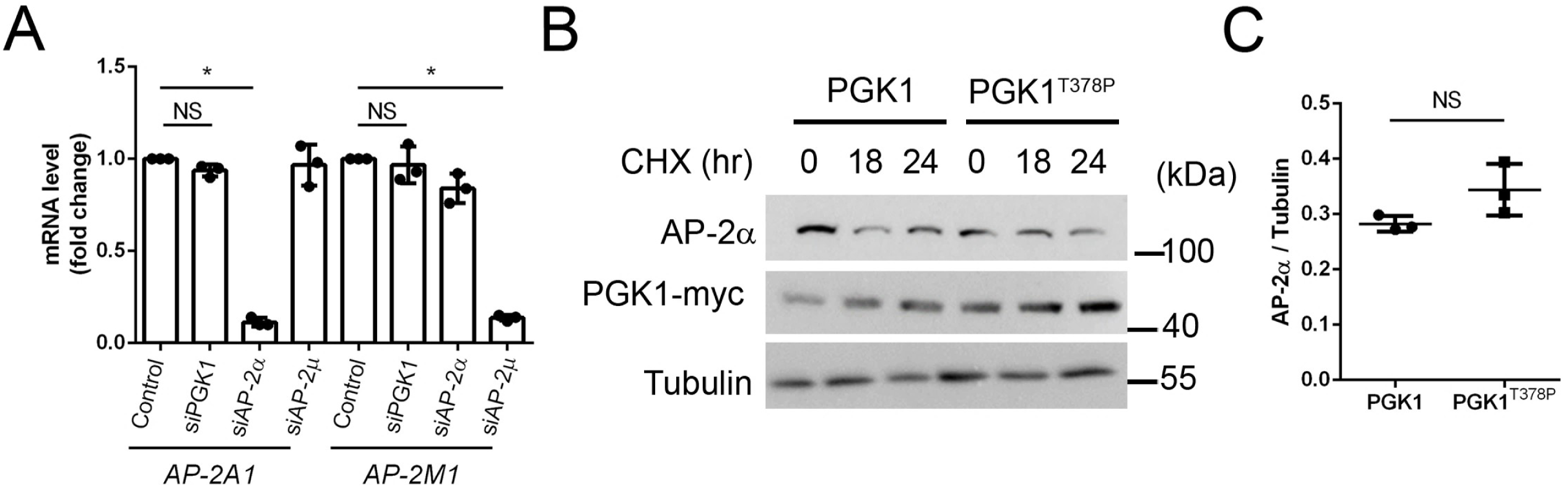
PGK1 affects the stability of AP-2 components. Data are shown as the means ± SDs. \**P*<0.05 NS *P*>0.05 **(A)** Total mRNA levels of *AP2A1* and *AP2M1*. One-way ANOVA with Tukey’s test: *AP2A1* – control vs. siPGK1 (*P*=0.4535), control vs. siAP-2α (*P*=2.09×10^-13^); *AP2M1* – control vs. siPGK1 (*P*=0.8624), control vs. siAP-2μ (*P*=9.38×10^-11^). **(B)** Time course analysis of AP-2α degradation upon CHX treatment to examine the effects of PGK1 and PGK1^T378P^. **(C)** Quantification of the AP-2α protein level normalized to the tubulin level at 24 hours (n=3). Paired Student’s t test, *P*=0.182.

**Figure EV4.**
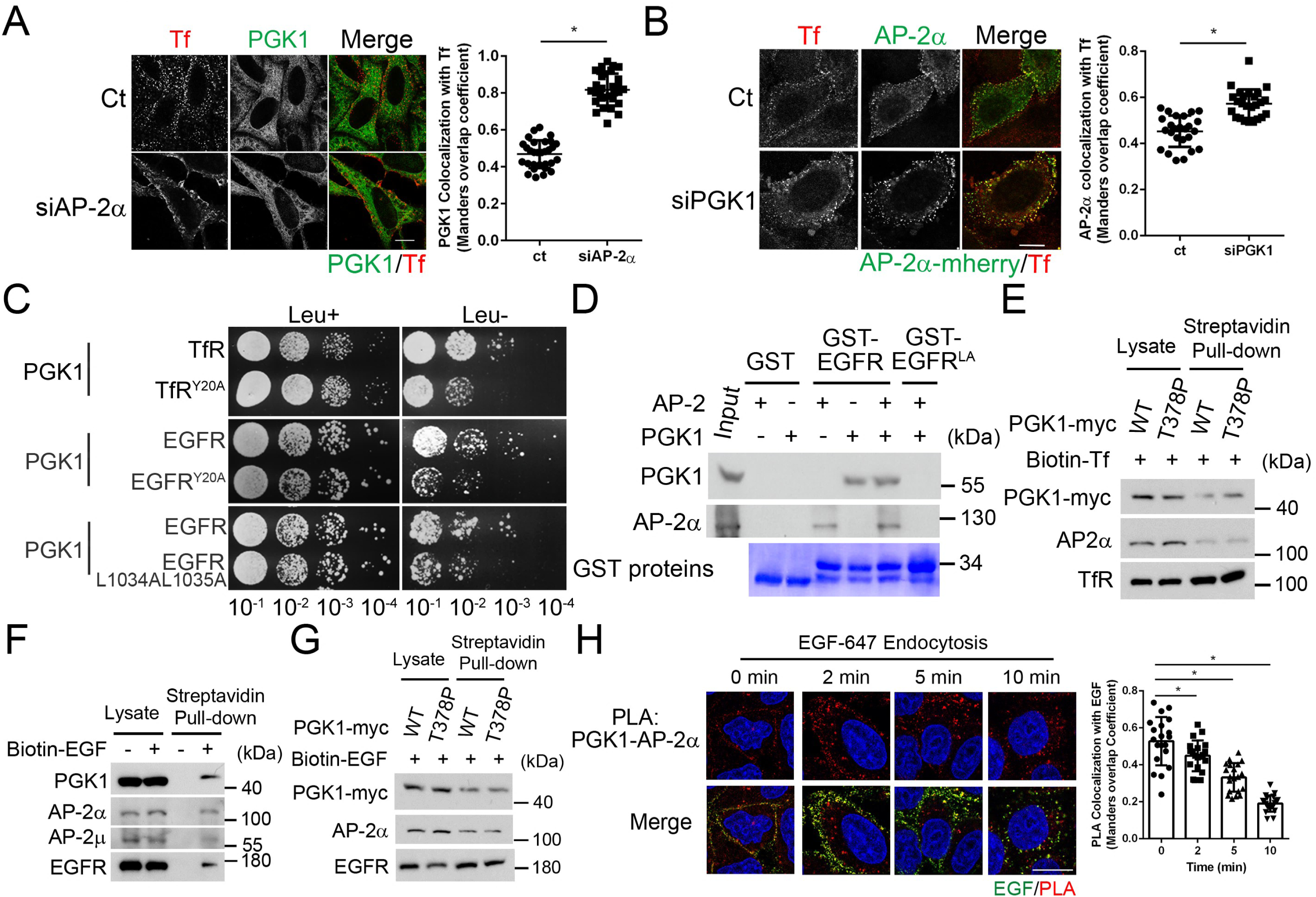
PGK1 recognizes cargo proteins to regulate endocytosis. Data are shown as the means ± SDs. \**P*<0.05 **(A)** Image (left; scale bar, 10 μm) and quantification (right) showing PGK1 (green) colocalized with Tf (red) in siAP-2α cells after 1 minute of internalization, expressed as the Manders’ coefficient (n=30 cells, two independent experiments). Unpaired Student’s t test, *P*=5.7×10^-24^. **(B)** Image (left; scale bar, 10 μm) and quantification (right) showing AP-2α-mCherry (green) colocalized with Tf (red) in siPGK1 cells after 1 minute of internalization, expressed as the Manders’ coefficient (n=25 cells, two independent experiments). Unpaired Student’s t test, *P*=4.5×10^-8^. **(C)** Yeast two-hybrid assay examining the interaction of PGK1 with sorting motifs on the cytoplasmic tail of TfR and EGFR and their mutations (n=3). **(D)** *In vitro* binding analysis of His-PGK1 and GST-EGFR fragments via GST pull-down to analyse the binding effect in the presence of AP-2α (n=3). **(E)** Myc-tagged PGK1 and PGK1^T378P^ associated with surface TfR and AP-2α (n=3). **(F)** Surface EGFR associated with endogenous PGK1 and AP-2 complexes (n=3). **(G)** Myc-tagged PGK1 and PGK1^T378P^ associated with surface EGFR and AP-2α (n=3). **(H)** PLA analysis showing colocalization of PGK1—AP-2α PLA puncta with EGF during EGFR endocytosis (n=20 cells, two independent experiments); scale bar, 10 μm. One-way ANOVA with Dunnett’s test vs. 0 min: 2 min (*P*=0.0187), 5 min (*P*=3.6×10^-9^); 10 min (*P*≈0).

**Figure EV5.**
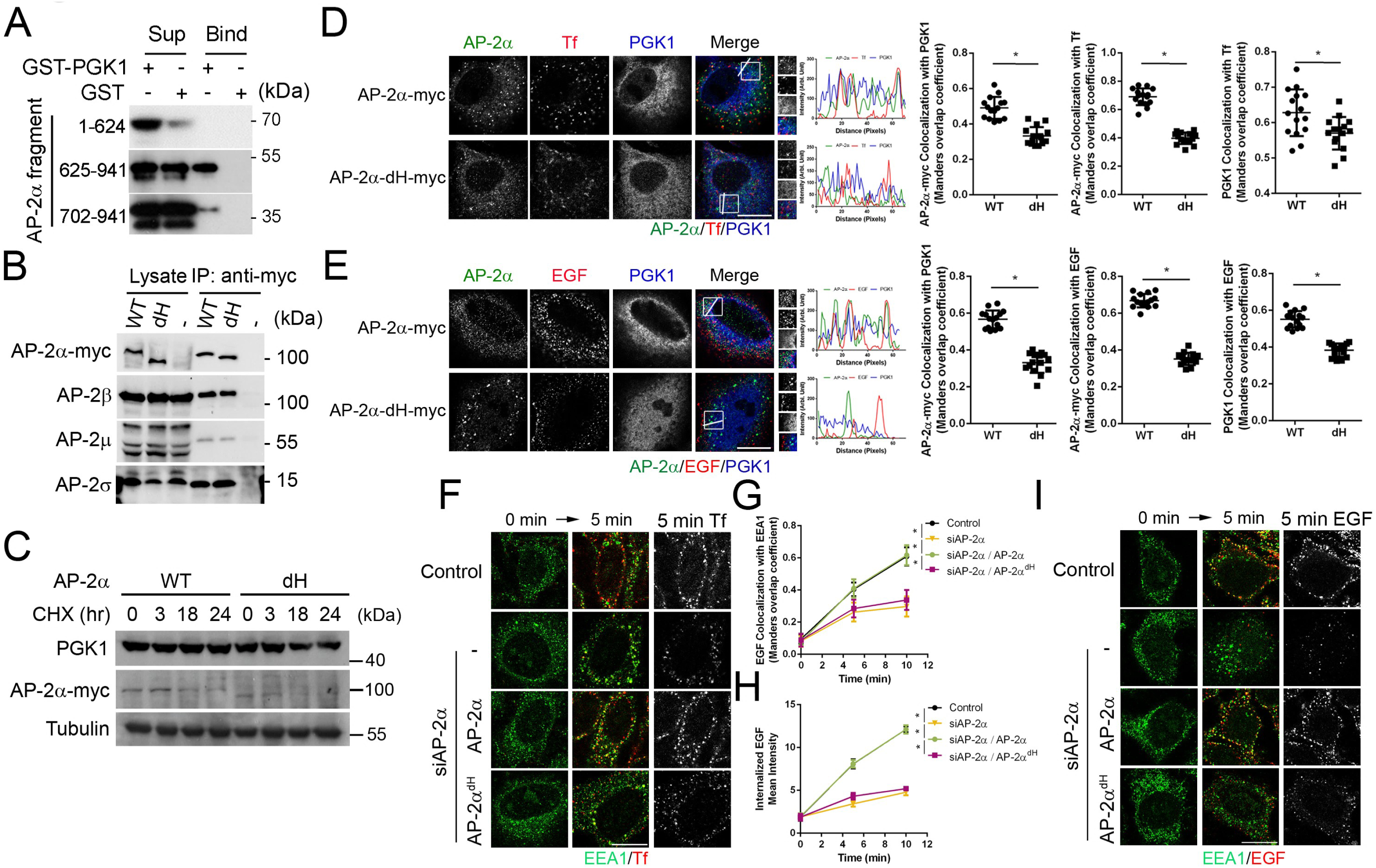
**The hinge region of AP-2**α **is required for its interaction with PGK1.** Data are shown as the means ± SDs. \**P*<0.05 **(A)** GST-PGK1 pull-down showing association with different His-tagged AP-2α fragments (n=3). **(B)** Immunoprecipitation of myc-tagged AP-2α and AP-2α^dH^ with AP-2 components in HEK293T cells (n=3). **(C)** Time course analysis of AP-2α degradation upon CHX treatment, examining AP-2α and AP- 2α^dH^ (n=3). **(D)** Confocal images (left; scale bar, 10 μm) and quantification (right), expressed as Manders’ coefficient, showing the effects of myc-tagged AP-2α and AP-2α^dH^ on colocalization of AP-2α- myc, Tf, and PGK1 during Tf internalization at 2 minutes (n=15 cells, two independent experiments). Unpaired Student’s t-test for WT vs. dH colocalization: AP-2α-myc with PGK (*P*=2.11×10^-8^), AP-2α-myc with Tf (*P*=1.81×10^-15^), and PGK1 with Tf (*P*=0.0091). **(E)** Confocal images (left; scale bar, 10 μm) and quantification (right), expressed as Manders’ coefficient, showing the effects of myc-tagged AP-2α and AP-2α^dH^ on colocalization of AP-2α- myc, EGF, and PGK1 during EGF internalization at 2 minutes (n=15 cells, two independent experiments). Unpaired Student’s t-test for WT vs. dH colocalization: AP-2α-myc with PGK (*P*=6.41×10^-12^), AP-2α-myc with EGF (*P*=1.48×10^-20^), and PGK1 with EGF (*P*=5.37×10^-13^). **(F)** Representative image of Tf (red) colocalized with EEA1 (green); scale bar, 10 μm. **(G)** Colocalization of EGF with EEA1, expressed as Manders’ coefficient, comparing the effects of AP-2α with AP-2α^dH^ in siAP-2α cells at 0, 5, and 10 minutes (n=15 cells, two independent experiments). One-way ANOVA with Tukey’s test at 5-minute time point: control vs. siAP- 2α (*P*=9.1×10^-9^); siAP-2α vs. AP-2α (*P*=2.8×10^-9^); AP-2α vs. AP-2α^dH^ (*P*=2.0×10^-7^). **(H)** Mean intensity of internalized Tf comparing the effects of AP-2α with AP-2α^dH^ in siAP- 2α cells at 0, 5, and 10 minutes (n=15 cells, two independent experiments). One-way ANOVA with Tukey’s test at 5-minute time point: control vs. siAP-2α (*P*=7.3×10^-12^); siAP-2α vs. AP- 2α (*P*=7.3×10^-12^); AP-2α vs. AP-2α^dH^ (*P*=7.3×10^-12^). **(I)** Representative image of EGF (red) colocalized with EEA1 (green); scale bar, 10 μm.

**Figure EV6.**
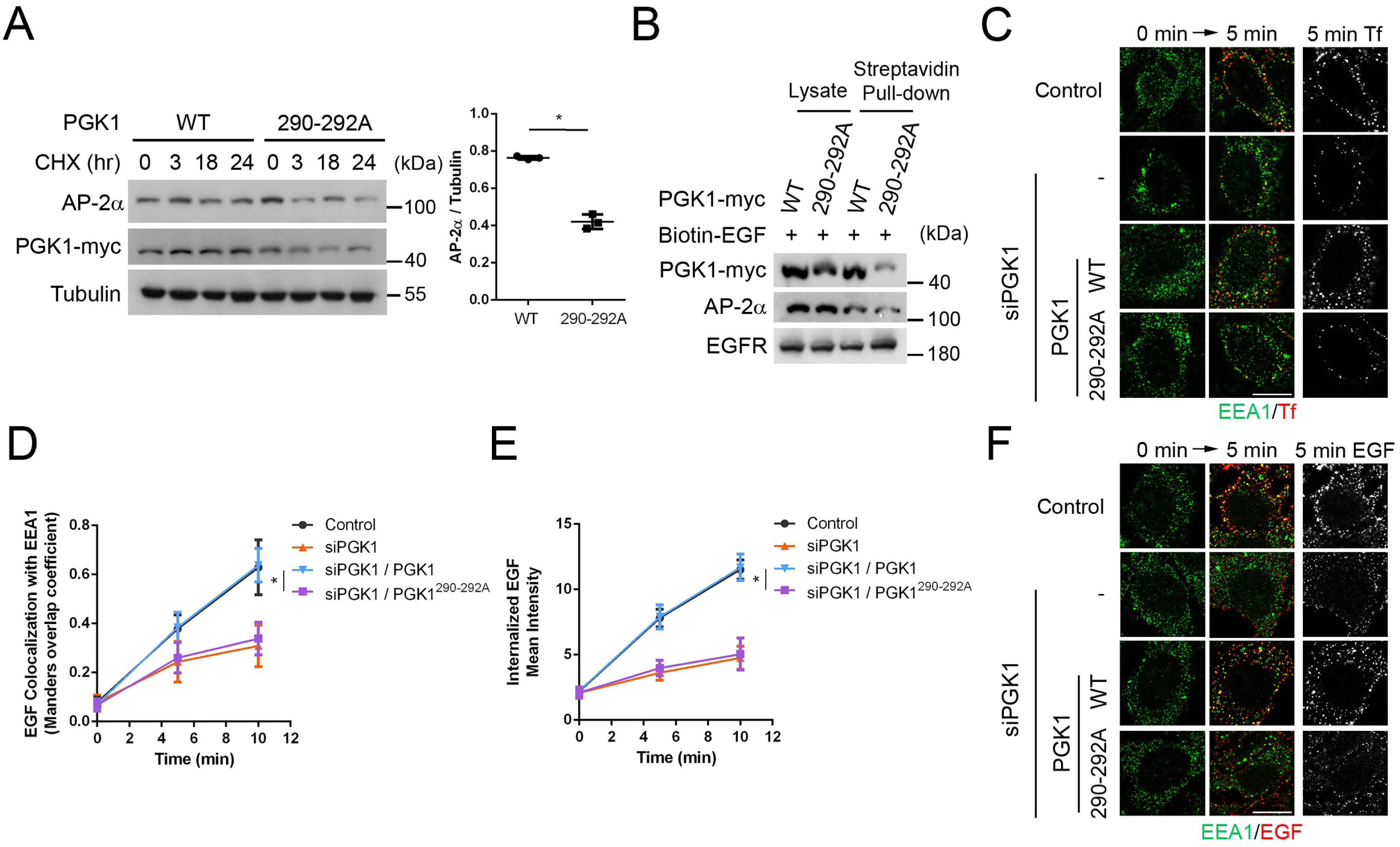
**AP-2**α **interacts with PGK1 via the DKF motif.** Data are shown as the means ± SDs. \**P*<0.05 **(A)** Time course analysis of AP-2α degradation upon CHX treatment, examining the effect of PGK1 and PGK1^290-292A^ (n=3). Quantification of the AP-2α protein level normalized to the tubulin level at 24 hours; Paired Student’s t test; *P*=0.0054. **(B)** Endogenous surface EGFR associated with endogenous AP-2α and myc-tagged PGK1 or PGK1^290-292A^ in siPGK1-treated cells (n=3). **(C)** Representative image of Tf (red) colocalized with EEA1 (green); scale bar, 10 μm. **(D)** Colocalization of EGF with EEA1, expressed as Manders’ coefficient, showing the effects of PGK1 and PGK1^290-292A^ in siPGK1 cells at 0, 5, and 10 minutes (n=15 cells, two independent experiments). Unpaired Student’s t-test at 5-minute time point for WT vs. 290-292, *P*=6.9×10^-6^. **(E)** Mean intensity of internalized Tf showing the effects of PGK1 and PGK1^290-292A^ in siPGK1 cells at 0, 5, and 10 minutes (n=15 cells, two independent experiments). Unpaired Student’s t-test at 5-minute time point for WT vs. 290-292, *P*=4.6×10^-14^. **(F)** Representative image of EGF (red) colocalized with EEA1 (green); scale bar, 10 μm.

**Figure EV7.**
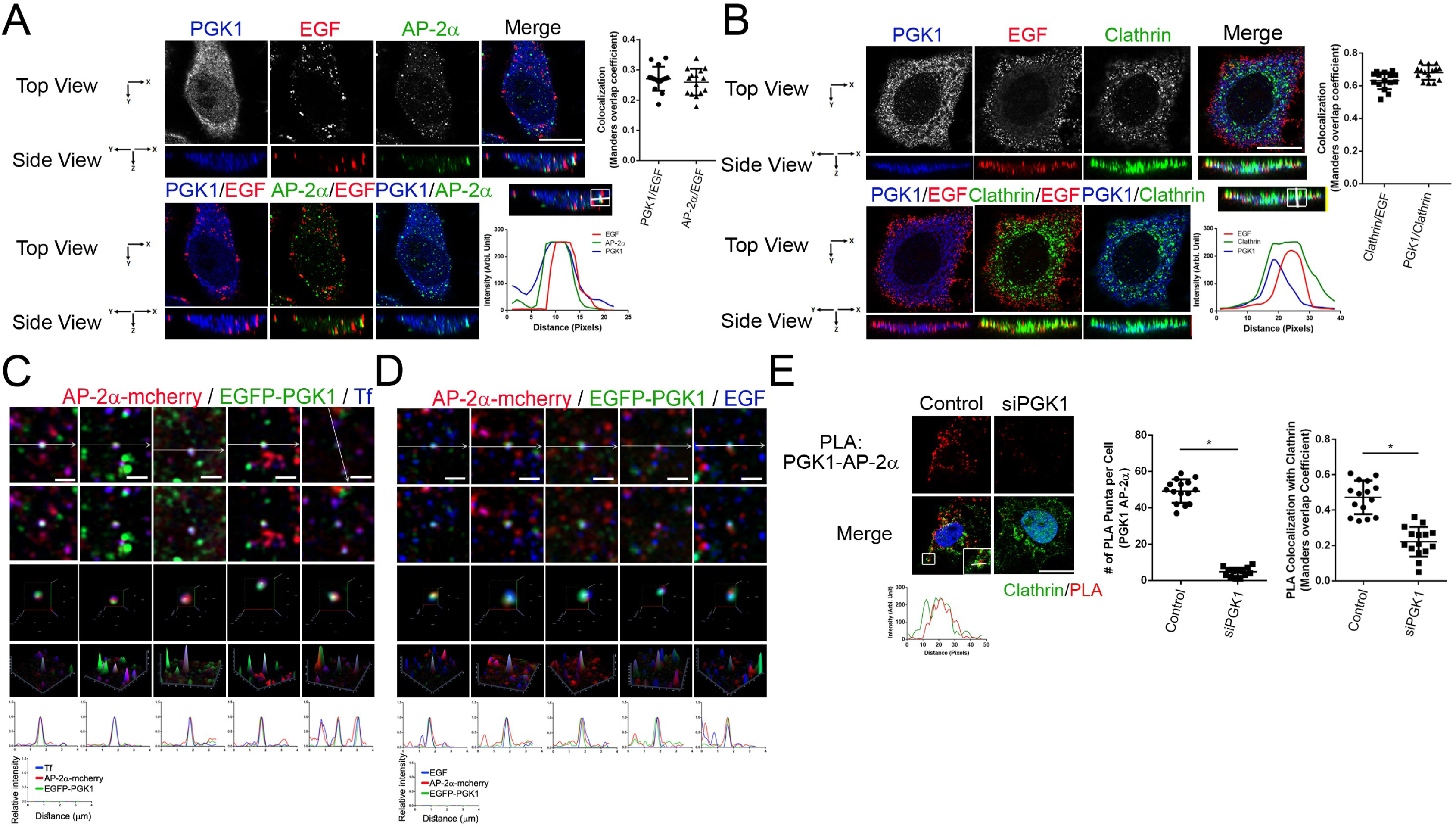
PGK1 colocalizes with clathrin and the AP-2 complex during protein endocytosis. Data are shown as the means ± SDs. \**P*<0.05 **(A)** Super-resolution confocal image showing colocalization of PGK1, EGF, and AP-2α after 2 minutes of internalization; scale bar, 10 μm. Quantification (right) of PGK1 with EGF and of AP- 2α with EGF colocalization, expressed as Manders’ coefficient (n=15 cells, three independent experiments). **(B)** Super-resolution confocal image showing colocalization of PGK1, EGF, and clathrin after 2 minutes of internalization; scale bar, 10 μm. Quantification (right) of PGK1 with clathrin and of clathrin with EGF colocalization, expressed as Manders’ coefficient; (n=15 cells, two independent experiments). **(C)** Colocalization analysis optimized by the prelysis method demonstrating that AP-2α-mCherry, EGFP-PGK1, and Tf colocalize in cells; scale bar, 1 μm. **(D)** Colocalization analysis optimized by the prelysis method demonstrating that AP-2α-mCherry, EGFP-PGK1, and EGF colocalize in cells; scale bar, 1 μm. **(E)** PLA analysis showing PGK1—AP-2α PLA puncta and Manders’ coefficient colocalization with clathrin (GFPspark-CTLB) (n=15 cells, two independent experiments); scale bar, 10 μm. Unpaired two-sided Student’s t test for control vs. siPGK1: puncta number (*P*=2.3×10^-8^); Manders’ (*P*=1.0×10^-20^).

**Figure EV8.**
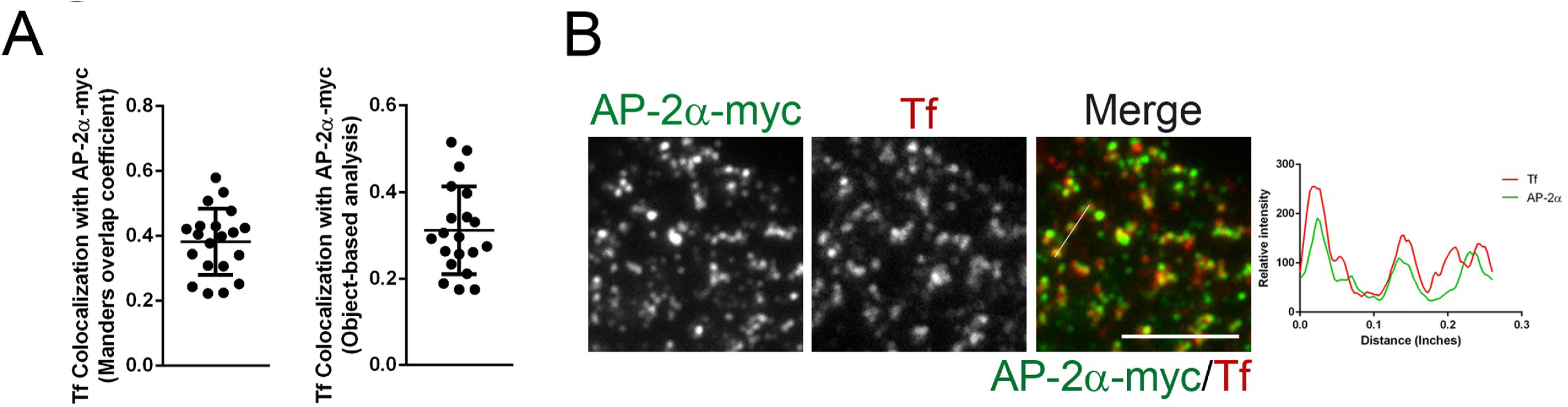
**TIRF imaging demonstrated that AP-2**α **recognizes Tf at the plasma membrane.** Data are shown as the means ± SDs. \**P*<0.05 **(A)** Quantitative results of Tf colocalization with AP-2α-myc, expressed in Manders’ coefficient (left) and object-based analysis (right) (n = 20 cells, two independent experiments). **(F)** TIRF imaging representing the colocalization of Tf and AP-2α-myc; scale bar, 5 μm. Arrows indicate the colocalization.

---

**Table EV1.** Proteins idetified from PGK1-interacting cell lysates.

| Accession number | Gene Name | Gene Description |
| --- | --- | --- |
| O75369 | FLNB | Filamin B |
| **O95782** | **AP2A1** | **Adaptor Related Protein Complex 2 Subunit Alpha 1** |
| P02533 | KRT14 | Keratin 14 |
| P02538 | KRT6A | Keratin 6A |
| P02768 | ALB | Albumin |
| P04259 | KRT6B | Keratin 6B |
| P04406 | GAPDH | Glyceraldehyde-3-Phosphate Dehydrogenase |
| P05783 | KRT18 | Keratin 18 |
| P05787 | KRT8 | Keratin 8 |
| P07355 | ANXA2 | Annexin A2 |
| P08195 | SLC3A2 | Solute Carrier Family 3 Member 2 |
| P08670 | VIM | Vimentin |
| P08729 | KRT7 | Keratin 7 |
| P08779 | KRT16 | Keratin 16 |
| P0CG38 | POTEI | Pote Ankyrin Domain Family Member I |
| P10809 | HSPD1 | heat shock protein family D (Hsp60) member 1 |
| P11021 | HSPA5 | heat shock protein family A (Hsp70) member 5 |
| P13645 | KRT10 | Keratin 10 |
| P13646 | KRT13 | Keratin 13 |
| P13647 | KRT5 | Keratin 5 |
| P14923 | JUP | Junction Plakoglobin |
| P15924 | DSP | Desmoplakin |
| P19013 | KRT4 | Keratin 4 |
| P19338 | NCL | Nucleolin |
| P21333 | FLNA | Filamin A |
| P22626 | HNRNPA2B1 | Heterogeneous Nuclear Ribonucleoprotein A2/B1 |
| P35908 | KRT2 | Keratin 2 |
| **P63010** | **AP2B1** | **Adaptor Related Protein Complex 2 Subunit Beta 1** |
| P68371 | TUBB4B | Tubulin Beta 4B Class Ivb |
| P78527 | PRKDC | Protein Kinase, Dna-Activated, Catalytic Subunit |
| P81605 | DCD | Dermcidin |
| Q01082 | SPTBN1 | Spectrin Beta, Non-Erythrocytic 1 |
| Q04695 | KRT17 | Keratin 17 |
| Q08380 | LGALS3BP | Galectin 3 Binding Protein |
| Q13813 | SPTAN1 | Spectrin Alpha, Non-Erythrocytic 1 |
| Q15149 | PLEC | Plectin |
| Q7Z794 | KRT77 | Keratin 77 |
| Q9BUJ2 | HNRNPUL1 | Heterogeneous Nuclear Ribonucleoprotein U Like 1 |
| Q9HBM0 | VEZT | Vezatin, Adherens Junctions Transmembrane Protein |
| Q9UJV9 | DDX41 | Dead-Box Helicase 41 |

